# High-Capacity Sample Multiplexing for Single Cell Chromatin Accessibility Profiling

**DOI:** 10.1101/2023.03.05.531201

**Authors:** Gregory T. Booth, Riza M. Daza, Sanjay R. Srivatsan, José L. McFaline-Figueroa, Rula Green Gladden, Scott N. Furlan, Jay Shendure, Cole Trapnell

**Affiliations:** Department of Genome Sciences, University of Washington, Seattle, WA, USA; Department of Biomedical Engineering, Columbia University, New York City, NY, USA; Brotman Baty Institute for Precision Medicine, Seattle, WA, USA; Allen Discovery Center for Cell Lineage Tracing, Seattle, WA, USA; Clinical Research Division, Fred Hutchinson Cancer Research Center, Seattle, WA, USA; Department of Pediatrics, University of Washington, Seattle, WA, USA; Howard Hughes Medical Institute, Seattle, WA, USA

## Abstract

Single-cell chromatin accessibility has emerged as a powerful means of understanding the epigenetic landscape of diverse tissues and cell types, but profiling cells from many independent specimens is challenging and costly. Here we describe a novel approach, sciPlex-ATAC-seq, which uses unmodified DNA oligos as sample-specific nuclear labels, enabling the concurrent profiling of chromatin accessibility within single nuclei from virtually unlimited specimens or experimental conditions. We first demonstrate our method with a chemical epigenomics screen, in which we identify drug-altered distal regulatory sites predictive of compound- and dose-dependent effects on transcription. We then analyze cell type-specific chromatin changes in PBMCs from multiple donors responding to synthetic and allogeneic immune stimulation. We quantify stimulation-altered immune cell compositions and isolate the unique effects of allogeneic stimulation on chromatin accessibility specific to T-lymphocytes. Finally, we observe that impaired global chromatin decondensation often coincides with chemical inhibition of allogeneic T-cell activation.

## INTRODUCTION

Millions of candidate regulatory DNA elements have been identified within mammalian genomes, many of which are specifically active in particular tissues and cell types(Thurman *et* al, 2012; Domcke *et al*, 2020; Cusanovich *et al*, 2018). Yet how these elements function, including how they respond to perturbation to modulate cells’ genetic programs, remains unclear. Moreover, given the importance of gene regulation in disease pathology, there is intense interest in therapeutic compounds targeting the epigenome or that modulate the activity of noncoding DNA(Kelly *et al*, 2010). However, many of these compounds target enzymes that contribute to the regulation of most if not all genes in the genome. Understanding the cell-type specific mechanisms of action of such compounds in healthy and diseased contexts is very challenging. With some epigenetic drugs already being applied in clinical settings, there is an urgent need for improved understanding of how epigenome-modulating compounds control the activity of our noncoding DNA(Bates, 2020; Kelly *et al*, 2010).

Sequencing based approaches which assess accessible regions of the genome have been fundamental for mapping the non-coding regulatory regions of the genome(Tsompana & Buck, 2014). Recent technological advances now enable assaying chromatin accessibility within single cells, thus resolving the chromatin states that comprise complex tissues and cell mixtures(Cusanovich *et al*, 2018; Satpathy *et al*, 2019). Thus far, technical advances have focused on increasing the number of cells profiled in a single experiment(Domcke *et al*, 2020; Lareau *et al*, 2019). However, it is often prohibitively expensive to perform these experiments on more than a few samples, with batch effects further complicating downstream analyses.

An increase in sample throughput would greatly broaden the applications for single cell genomic technologies. For instance, high throughput chemical screens (HTS) have been foundational in identifying candidate compounds which mitigate disease(Broach & Thorner, 1996; Pereira & Williams, 2007). Pairing HTSs with measurements of global gene expression in bulk(Bush *et al*, 2017; Ye *et al*, 2018), or single cells(McGinnis *et al*, 2019; Stoeckius *et al*, 2017; Shin *et al*, 2019; Gehring *et al*, 2020; Srivatsan *et al*, 2020), has improved the resolution at which one can evaluate therapeutic responses. Furthermore, the ability to resolve molecular phenomena within single cells will be important in understanding the heterogeneous response to therapy seen within and between individuals and identifying the molecular determinants of therapeutic resistance(Shaffer *et al*, 2017)-(Mathur & Roberts, 2018; Alfert *et al*, 2019).

We recently introduced sciPlex, an inexpensive and robust strategy for multiplexing combinatorial indexing-based single-cell RNA-seq experiments(Srivatsan *et al*, 2020). This strategy (to which we refer here as ‘sciPlex-RNA-seq’) exploits a propensity for permeabilized nuclei to absorb unmodified single stranded DNA oligos of any sequence, referred to as hash labels. During sci-Plex-RNA-seq, permeabilized cells or nuclei from distinct samples are incubated with unique hash labels (or combinations thereof), which are then stabilized within nuclei through chemical fixation. These cells or nuclei are then sequenced using sci-RNA-seq(Cao *et al*, 2017, 2019), which recovers the hash labels for each cell along with its transcriptome.

Here, we extend the sciPlex multiplexing platform to combinatorial indexing-based chromatin accessibility profiling within single cells. We refer to this approach as sciPlex-ATAC-seq (**Fig. 1a**). Using this nuclear labeling scheme, we first resolve treatments of cells from a pooled chemical screen and pair this information with their chromatin accessibility profiles. We further adapt this multiplexing strategy for highly scalable single cell accessibility profiling, which we apply to human mixed lymphocyte reactions (MLR) from multiple donors in combination with chemical perturbation. From these pooled assays we identify allogeneic-dependent chromatin changes within activated T-lymphocytes, which are disrupted by immunosuppressive compounds.

**Fig. 1:**
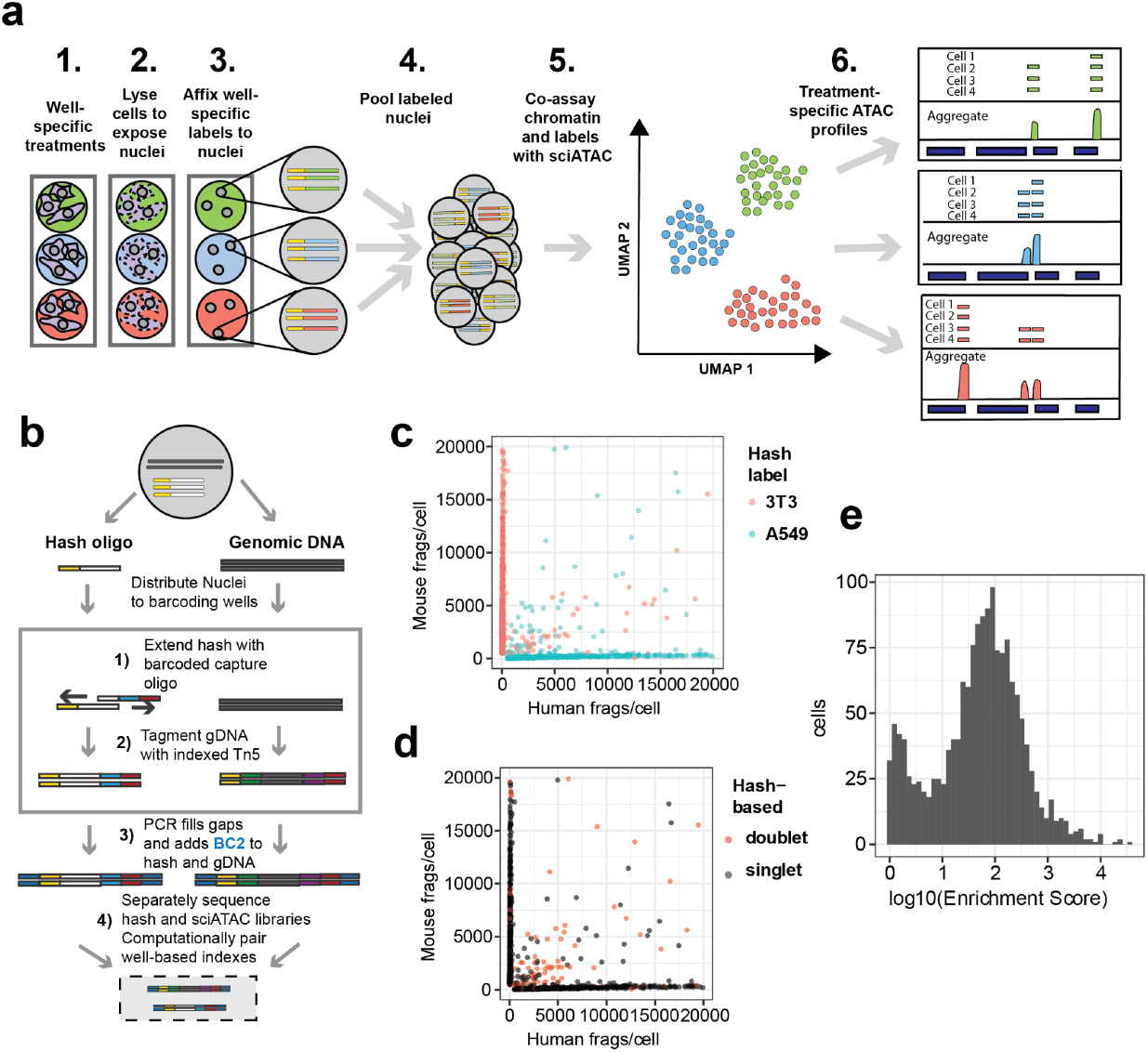
Single stranded DNA oligos label nuclei enabling sample multiplexing single nucleus chromatin accessibility profiling. **a)** Experimental workflow. **b)** Schematic of co-assay strategy for label capture and chromatin accessibility profiling within individual nuclei through two-level combinatorial indexing. **c)** Scatter plot of mouse and human unique fragment counts for individual cells colored by co-assayed nuclear hash labels for each nucleus. **d)** Scatter plot of mouse and human unique fragment counts colored by doublet calls based on co-assayed hash labels. **e)** Histogram of hash label enrichment scores for each nucleus, where *Enrichment Score = x/y*, where *x* = counts for the most common hash label within a cell and *y* = the second most common hash label within a cell. For cells with only one hash ID, the enrichment score was set as the number of hash reads (to avoid infinite values).

## RESULTS

### Hash labeling enables multiplexing of sciATAC-seq samples

To capture poly-adenylated hash oligonucleotides within the context of sciATAC-seq, we modified the original sciATAC-seq protocol(Cusanovich *et al*, 2015), to include an indexed primer extension step. The indexed extension of the hash oligos is followed by indexed transposition within the same well, creating a known pairing between well barcodes of hash oligos and tagmented chromatin. By design, extended hash oligos and tagmented chromatin possess Nextera P5 and P7 PCR handles. After tagmentation, nuclei are pooled, stained with DAPI, and flow-sorted into 96 well plates for crosslink reversal and PCR. Amplification with indexed PCR primers adds an additional level of barcoding to both hash labels and accessible chromatin fragments. In sum, this strategy leverages unique combinations of well barcodes on hash and chromatin fragments to pair single cell chromatin profiles with their corresponding hash identifiers (**Fig. 1b**).

To determine the specificity of hash labeling, we performed a species mixing experiment. A roughly equal number of freshly expanded NIH-3T3 cells (mouse) and A549 cells (human) were distributed to distinct wells of a 96-well v-bottom plate, where they were permeabilized and each species affixed with a different set of hashing oligos (**Supp. Fig. 1a**). Upon library preparation and sequencing, we were able to accurately identify the species of each recovered nuclei based on the top enriched hash label recovered (99%, n = 1696) (**Fig. 1c**). Moreover, half of the barcode collision events observed, which are identified by a mixture of human and mouse chromatin (doublets, etc.), could be readily identified by the presence of more than one enriched hash label (n = 127) (**Fig. 1d**). Moreover, a proportion of the doublets resulting from a barcode collision between two or more nuclei of the same species, were identified using the exogenous hash labels (n = 86).

By recovering many uniquely indexed hash molecules per nucleus, we can quantitatively assess label mixing across sciPlex experiments. Hash enrichment scores, which reflect the ratio of the most abundant label to the second most abundant molecule per recovered nucleus, revealed a 100-fold enrichment of top hashes on average, suggesting minimal label diffusion between nuclei during library preparation (**Fig. 1e**). In an effort to identify optimal conditions for sciPlex-ATAC-seq, we examined various permeabilization, hash-extension, and tagmentation conditions, all of which produced comparable quality sciATAC libraries and hash labeling (**Supp. Fig. 1a-d**). We also considered whether the fixation process might lead to increased intra-sample multiplets. By performing species mixing experiments where each cell line was either hashed and fixed separately (post-mix), or hashed and fixed after combining the cell lines (pre-mix) we were able to compare the final observed collision rates. While the pre-mix sample did produce a higher observed collision rate (pre = 23.9%, post = 14.7%), the collision rate was similar to the expectation for this experiment (20%).

### Multiplexed single-cell ATAC profiles reveal drug-specific and dose-dependent changes in the chromatin landscape

The ability to pool many samples for parallel processing should reduce the potential for batch-to-batch variation, while simplifying the handling required for experiments with numerous conditions and/or replicates. We therefore sought to apply sciPlex-ATAC-seq to resolve chromatin profiles at single cell resolution in the context of a multi-compound chemical perturbation experiment.

For this chemical screen, we mirrored chemical perturbations we previously found to elicit diverse and dose-dependent effects on cells(Srivatsan *et al*, 2020). Selected compounds included Dexamethasone (Dex), a glucocorticoid receptor agonist, Vorinostat (SAHA), a broad spectrum histone deacetylase inhibitor, Nutlin-3A, an MDM2 inhibitor, which increases P53 activity(Wade *et al*, 2013), and BMS-345541 (BMS), an inhibitor of NFkB-dependent transcription. Human lung adenocarcinoma-derived cells (A549) were cultured in a 96-well dish and treated for 24 hours with one of the four compounds at eight different concentrations covering three orders of magnitude, including vehicle control only. Each condition was also performed in triplicate (**Fig. 2a**). After removing low quality profiles (**Supp. Fig. 3a**), we assigned treatment labels to all cells with at least 10 hash counts and a minimum of 2-fold label enrichment (**Supp. Fig. 3c**,**d**). In addition to removing doublets based on poor hash labeling, we applied a modified version of scrublet(Wolock *et al*, 2019; Domcke *et al*, 2020) to identify and prune any remaining doublets (**Supp. Fig. 3b**). After filtering, we recovered a total of 8,655 cells. The number of cells recovered was strongly related to the dosage of each compound, with the exception of Dex (**Supp. Fig. 4a**). Furthermore, based solely on the number of filtered cells recovered per condition, we were able to derive kill curves and estimate IC50 values for each compound, similar to those derived previously (**Supp. Fig. 5**)(Srivatsan *et al*, 2020). While per-cell statistics such as hash recovery and total fragments per cell were largely consistent across conditions (**Supp. Fig. 4b, d**), regressing TSS enrichment of chromatin fragments as a function of dose revealed a significant reduction upon exposure to BMS (coef = -0.36; p = 1.13e-48) and SAHA (coef = -0.14; p = 1.47e-13), indicating a more dispersed distribution of transposase insertion sites in the presence of these drugs (**Supp. Fig. 4c**).

**Fig. 2:**
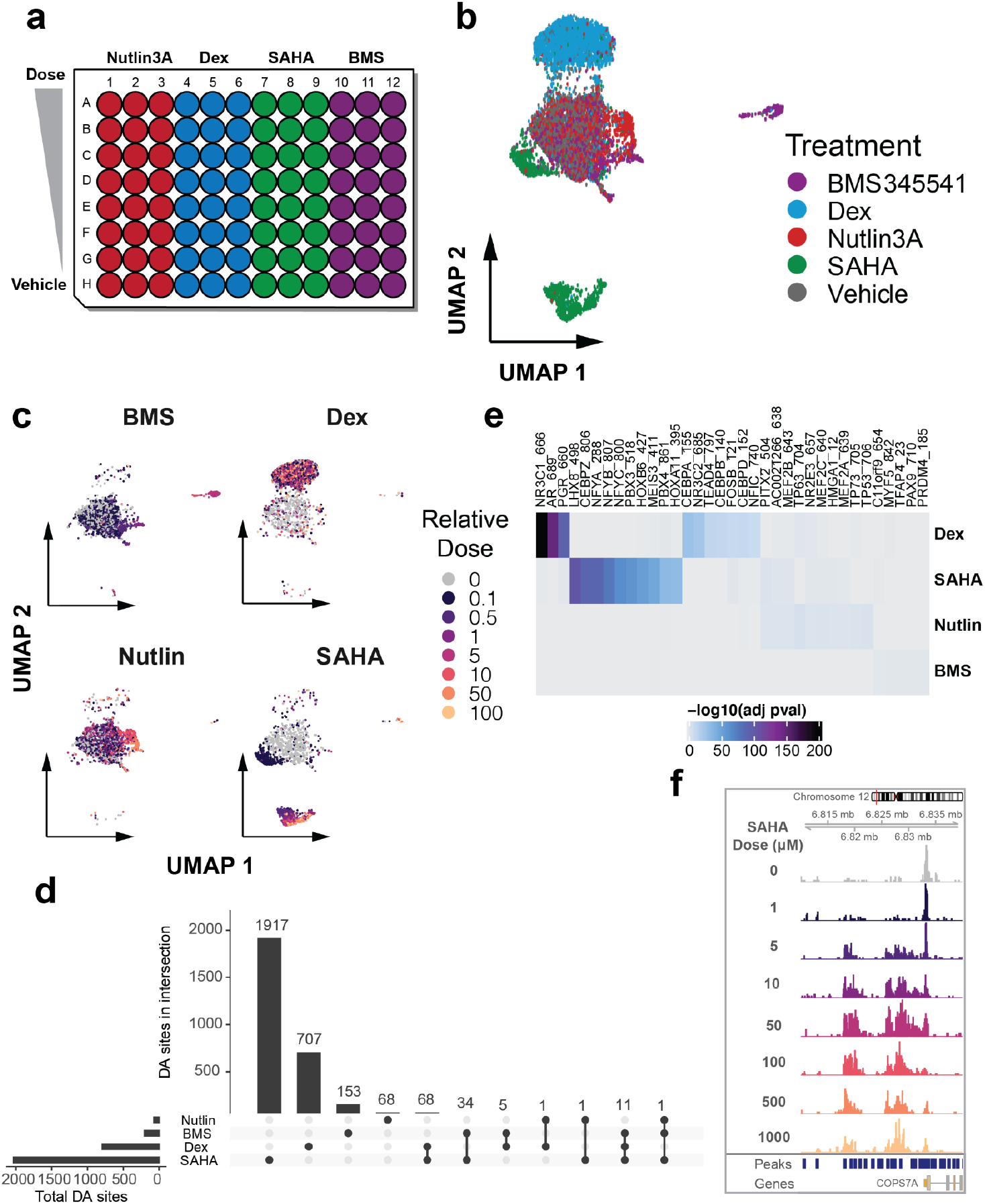
Nuclear hashing enables scalable single-cell epigenomics for high throughput chemical screens. **a)** Schematic of chemical screen culture dish layout. **b)** UMAP embedding of single cell chromatin profiles colored by drug treatment group labels. **c)** Faceted UMAP embeddings of cells from each treatment group colored by relative dose labels. **d)** Upset plot displaying the total number of DA sites identified in response to each drug and numbers of shared DA sites across treatment groups. **e)** Heatmaps depicting up to 10 most significantly enriched motifs per drug within peaks found to open with treatment. **f)** Browser tracks for cells aggregated by dose of the SAHA compound. The y-axis represents read coverage ranging from 0 - 20 (normalized by reads within promoters for each group.

Upon dimensionality reduction of the scATAC profiles, cells reproducibly inhabited chromatin states largely defined by the drug treatment received (**Fig. 2b, Supp. Fig. 6a**). Moreover, by labeling cells by the dosage of each drug exposure, we could investigate how the concentration of each compound affects the chromatin landscape. Cells treated with BMS showed chromatin states that abruptly diverged from vehicle treated cells at doses exceeding 1uM, possibly reflecting toxicity-induced effects. A Dex-induced chromatin state was attained even at the lowest concentrations examined, suggesting a more binary and stable impact of the corticosteroid mimic. And while Nutlin-3A induced very few detectable changes, SAHA treated cells exhibited a severe and dose-dependent progression away from the vehicle-treated state (**Fig. 2c**).

Using clusters of various chromatin states identified with Monocle3(Cao *et al*, 2019), we called de-novo peaks of accessibility, identifying a total of 129,410 accessible sites across all conditions in our chemical screen (**Supp. Fig. 6b**,**c**). To identify sites in our experiment with accessibility impacted by each chemical perturbation, we modeled the accessibility of each peak as a function of drug dose. We identified 2966 sites across the genome whose accessibility was significantly impacted by a chemical perturbation in our experiment (promoters = 301, distal = 1232, intronic = 1223, exonic = 210) (**Supp. Table S1**). Interestingly, the vast majority of differentially accessible sites were specific to just one compound, with the HDACi SAHA altering the most sites (opening = 890, closing = 1142) (**Fig. 2d**).

To identify the sites that may be directly bound by the transcription factors (TFs) targeted by these compounds, we assessed the enrichment of TF binding elements within sites that significantly gained or lost accessibility in response to each compound (**Fig. 2e** and **Supp. Fig. 7**). Confirming the known effect of dexamethasone in inducing nuclear localization and DNA binding of the glucocorticoid receptor (GR)(McDowell *et al*, 2018), we found the binding site for GR (NR3C1) to be the most enriched motif within sites opening in response to Dex treatment (**Fig. 2e**). Notably, binding sites for androgen- (AR) and progesterone receptors (PGR) were also highly enriched in Dex-opening sites, likely owing to target sequence similarities with GR(Nelson *et al*, 1999). Despite a limited global impact of Nutlin-3A on chromatin accessibility, the binding site for P53 (TP53) and homologs P63 and P73 were all among the top five most enriched elements within Nutlin-3A induced accessible sites, consistent with the drug’s role in activating P53 (**Fig. 2e**). Visually inspecting the accessibility data grouped by treatment at significantly affected loci, supported the altered state of chromatin for each treatment (**Fig. 2f, Supp. Fig. 8**). Together, these results point to unique and discernible effects of each tested compound on the chromatin landscape in ways which reflect the activity of targeted factors.

### Dose-dependent changes in the chromatin landscape reflect the transcriptional state of individual cells

We next applied an unsupervised, graph learning approach (Methods)(Cao *et* al, 2019; Qiu *et al*, 2017), to identify cells’ trajectories through altered chromatin states as a function of the dose of each drug (**Fig. 3a, Supp. Fig.8a**,**d**,**g**). For each cell we defined a “pseudodose” as its position within the identified trajectory (**Fig. 3a-c, Supp. Fig. 8a-i**). Cells treated with dexamethasone again exhibited one of two discrete chromatin states, with all tested doses of Dex producing similar changes in chromatin accessibility (**Supp. Fig. 9a-c**). Across the Dex pseudodose axis, significantly Dex-affected regions were largely found to exhibit increasing (n = 312) or dynamic (n = 367) accessibility (decreasing; n = 109) (**Supp. Fig. 10a**,**d**), with increasingly accessible promoters and distal elements enriched for GR (NR3C1) motifs (**Supp. Fig. 10b**,**c**).

**Fig. 3:**
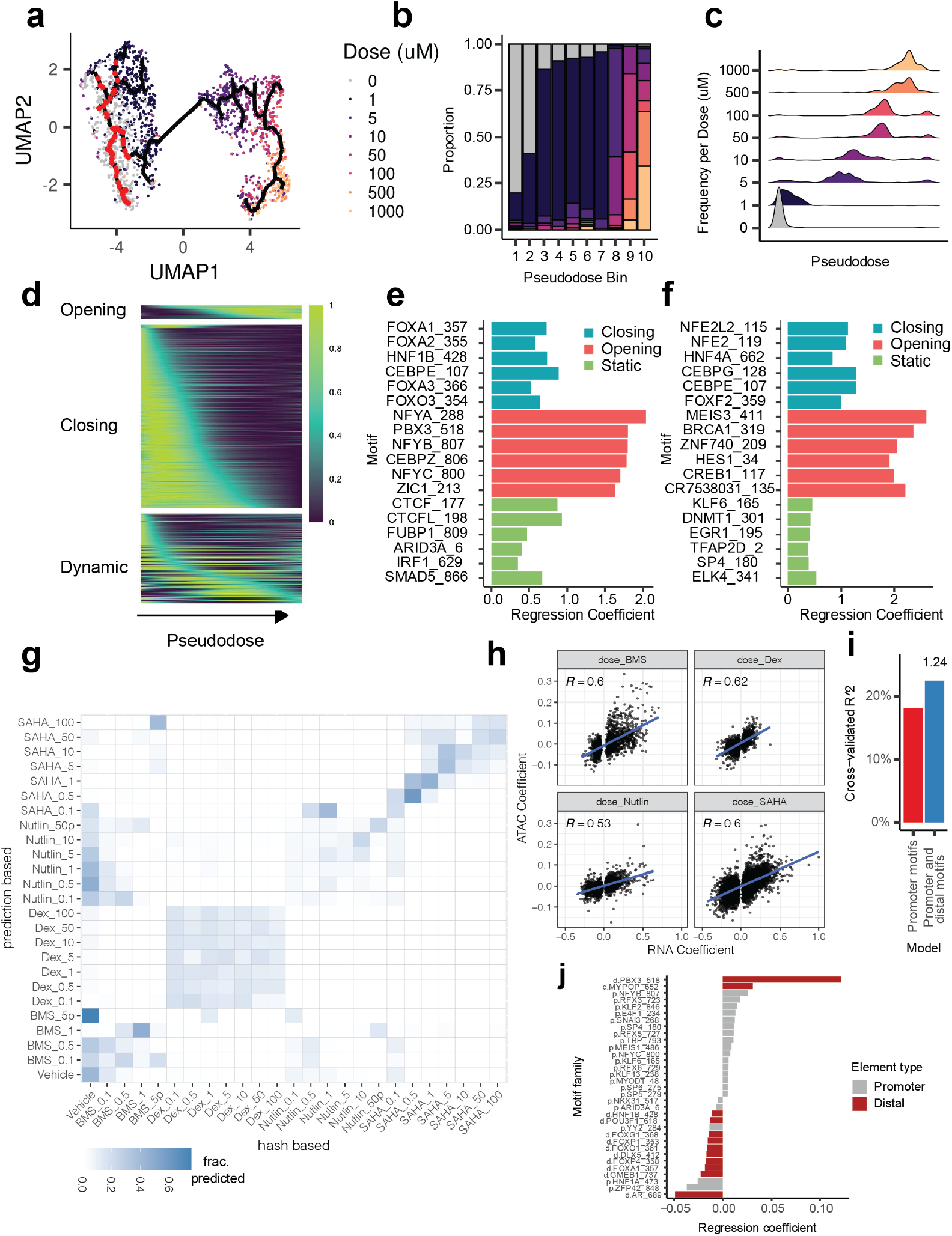
HDACi treatment induces dose-dependent changes in the chromatin landscape, which reflect the transcriptional state of individual cells. **a)** UMAP of vehicle and SAHA-treated cells showing the predicted chromatin state trajectory through which cells traverse with increasing doses of SAHA. Data points are colored by doses used. Red points on trajectory represent ‘root’ positions (located within groups of vehicle-treated cells). **b)** Proportions of cells treated with each actual dose of SAHA across pseudodose trajectory bins. **c)** Density plots quantitate trajectory positions of cells treated with increasing doses of SAHA. **d)** Smoothed accessibility scores across SAHA-pseudodose for three classes of identified differentially accessible sites (opening, closing, and dynamic). Closing and opening sites were defined as sites with a maximum accessibility score occurring within the first or last 20 pseudotime bins (out of 100 bins), respectively. Dynamic sites have a maximum score in the intervening pseudodose bins. **e & f)** Top motifs explaining whether a distal (E) or promoter-proximal (F) SAHA-DA site is classified as closing (blue), opening (red) or static (non-DA, green). **G**. Treatment label prediction accuracy when mapping sciPlex-ATAC data to corresponding sciPlex-RNA (Srivatsan *et al*, 2020) datasets. Labels from scRNA-seq cells to our cells with matched scATAC information using Seurat’s find anchor transfer function (Stuart *et al*, 2019), implemented by the ArchR package (Granja *et al*, 2021). Cells which received BMS dose treatments greater than 1*µ*M were pooled and labeled BMS_5P due to very low cell recovery for these conditions. **H**. Scatter plots comparing regression coefficients for gene accessibility scores (ATAC) and gene expression (RNA) as a function of dose for each drug. Only genes with significant dose terms for gene expression were considered. **I**. SAHA-responsive DE Gene expression variation explained by models taking into account sequence elements within promoters alone, or within promoters and distal co-accessible sites. The value (1.24) above the right bar reflects the fold increase in predictive power with models which include distal co-accessible sites. **J**. Sequence elements with largest regression coefficients from promoter (grey) or distally connected sites (red) when predicting SAHA-affected gene responses.

We hypothesized that a cell’s chromatin state could be used to infer impacts of each treatment on the transcriptome. Here we used accessibility scores for each gene (Methods) in order to integrate our single cell chromatin profiles with previously generated single cell transcriptomes from an identical set of chemical perturbations(Srivatsan *et al*, 2020). After integrating the matched data sets, we transferred treatment labels from the nearest single cell transcriptomes in shared, reduced dimension space, onto our single cell chromatin profiles(Stuart *et al*, 2019) (Methods). Transferred labels from associated transcriptomes consistently reflected the true compounds driving the chromatin profiles (**Fig. 3g**). Strikingly, the dosages of Dex treatments were often misidentified, further emphasizing the dose-independent response of cells to Dex within the tested range. Nonetheless, gene-specific impacts were highly correlated at the chromatin and transcript level across compounds (**Fig. 3h**).

Ultimately, we sought to identify sequences in noncoding DNA that would predict transcriptional responses of nearby genes to Dex. Applying co-accessibility scores(Pliner *et al*, 2018) to link promoters with distal elements, we used sequence features to generate predictive models of drug-induced gene expression changes (Methods). Including TF motifs within distal sites that were linked to Dex-affected gene promoters improved predictions of gene expression by more than 50%, compared with models based on promoter elements alone (**Supp. Fig. 10e**). Reassuringly, distally located GR binding motifs had the largest effect size in predicting whether genes increase expression in response to Dex-treatment (**Supp. Fig. 10f**).

The effects of BMS or Nutlin-3A on gene accessibility were also correlated with transcript abundance (Fig. 3h). However, because of the more modest impact of these compounds on both chromatin and transcript abundance, BMS and Nutlin-3A treated profiles were often classified as untreated and vice versa. (Fig. 3g). With comparatively few significant changes in accessible regions from BMS and Nutlin-3A treatments, further analysis of their relationship to gene expression was limited.

The histone deacetylase inhibitor SAHA produced the clearest dose-dependent trajectory of alterations in chromatin accessibility. Grouping SAHA-treated cells by pseudodose values revealed a strong correspondence between trajectory position and the true treatment dose, suggesting that the trajectory reflects increasingly SAHA-altered chromatin state (**Fig. 3b**). Interestingly, at intermediate SAHA doses (5 - 100 *µ*M), individual cells occupied more broad pseudodose ranges, indicative of a heterogeneous progression towards the maximally SAHA-affected state (**Fig. 3c**). Counterintuitively, more SAHA-affected sites exhibited a sustained loss (n = 757), rather than a gain (n = 58), of accessibility along the pseudodose axis (**Fig. 3d, Supp. Fig. 11**), a trend that was poorly explained by specific TF binding motifs (**Fig. 3e** and **3f**). A dose-dependent reduction of accessibility in HDACi-treated cells has been observed previously(Sanchez *et al*, 2018; Pattenden *et al*, 2016), and together these results support a general requirement for the turnover of histone acetylation in maintaining proper nucleosome organization.

The impacts of SAHA on expression and accessibility were highly correlated for affected genes (**Fig. 3h**). Moreover, shared, dose-specific effects on both chromatin and RNA abundance of SAHA-treated cells enabled accurate dosage-level treatment predictions for individual chromatin profiles (**Fig. 3g**). Similar to the Dex response, when predicting the effect of SAHA exposure on a gene’s expression, including sequence features within distally connected sites improved predictive power (**Fig. 3i**). The motif for PBX3, which is enriched within SAHA-opened sites (**Fig. 2e, Fig. 3e**), was also the strongest predictor for increased gene expression upon SAHA treatment when present within distal sites co-accessible with a gene’s promoter (**Fig. 3j**). Together these data warrant further investigation of a role for PBX3 in the HDACi response. Ultimately, when applied to HTS experiments, sciPlex-ATAC-seq highlights the underlying contribution of chromatin, often within non-coding regions, in governing cellular response to chemical perturbation.

### Improved scalability enables donor and cell-type specific analysis of chromatin organization in mixed lymphocyte reactions

Investigating heterogeneous cell mixtures from each of many conditions necessitates a platform which easily scales beyond 10,000-100,000 cells. We therefore adapted our nuclear hashing approach for compatibility with the recently described three-level combinatorial indexing strategy for highly-scalable single cell chromatin accessibility profiling(Domcke *et al*, 2020). We refer to this adapted protocol as sciPlex-ATAC3.

For sciPlex-ATAC3, accessible chromatin is tagged using a single commercially-available transposome complex. Two separate splinted ligation reactions are then used to sequentially add well-specific barcodes separately to each side of the transposed DNA fragments, followed by PCR-based barcode addition. We reasoned that with hash oligos bearing the universal Nextera P7 sequence and a capture oligo possessing the Nextera P5 sequence, the resulting annealed product would resemble transposed chromatin fragments and be susceptible to the same indexing reactions as chromatin within each nucleus (**Supp. Fig. 12**). We evaluated sciPlex-ATAC3 with species-mixing experiments and found that by increasing the melting temperature of the annealed hash-capture product, we were able to achieve accurate label assignment without the need for a hash extension step (**Supp. Fig. 13**).

Using human peripheral blood mononuclear cells (PBMCs) for mixed lymphocyte reactions as an experimental system, we sought to apply sciPlex-ATAC3 to explore allogeneic immune responses across multiple donors. PBMCs were harvested from four unrelated individuals and split into two fractions, responders and stimulators, the latter of which were subsequently irradiated. The two fractions were recombined such that cells from each responder were paired 1:1 with stimulators from all other donors (and autologously as a control) and cultured in 96-well plates. Additional controls included responders cultured with stimulatory beads (ratio 10:1; responder : bead), and both responders and stimulators cultured alone (**Fig. 4a**). Within the 96-well culture plate, each of the 28 conditions was cultured in triplicate for five days prior to harvesting for sciPlex-ATAC3 or flow cytometric analysis (**Supp. Fig. 14a**). Nuclei from each well were then hashed and pooled for sciPlex-ATAC3, allowing us to assay chromatin profiles from single nuclei, simultaneously for all conditions and biological replicates.

**Fig. 4:**
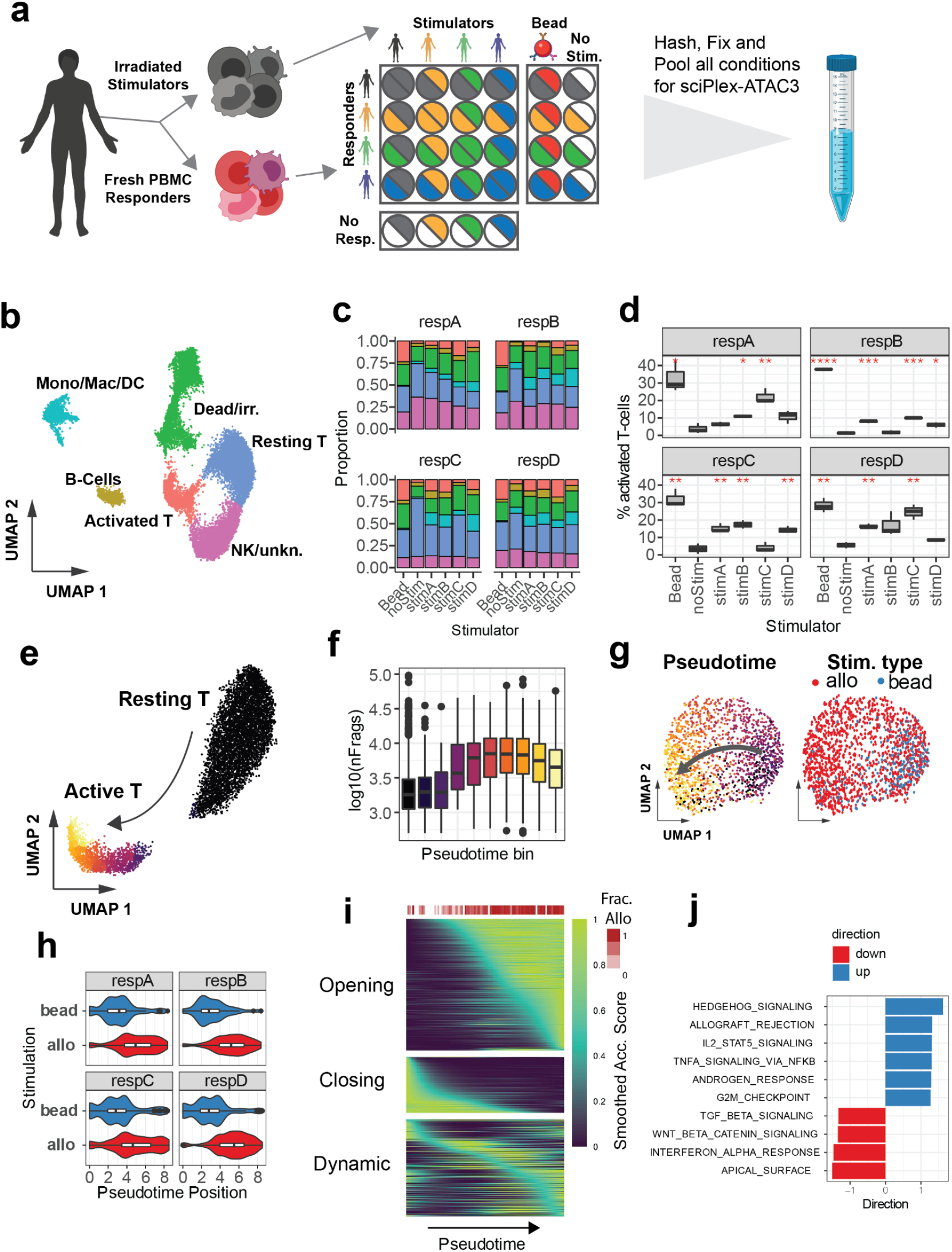
sciPlex-ATAC3 elucidates effects of allogeneic T-cell stimulation on the chromatin landscape. **a)** Schematic of mixed lymphocyte reaction conditions and experimental setup. **b)** UMAP representation of chromatin profiles from recovered cells with assigned cell type annotations. **c)** Proportion of cells across annotations for each experimental condition (excluding the “stimulator alone” condition). **d)** Percentage of activated T-cells recovered from each condition. Variation in recovery was determined by separately quantifying each of three biological replicates for all conditions. Red asterisks indicate significant difference in the mean relative to no-stim for each responder (Student’s t-test; *: p <0.05; **: p <0.01, ***: p < 0.001; ****: p < 0.0001). **e)** UMAP representation of chromatin profiles of T-cells from the MLR experiment, colored by pseudotime bins as determined using the *learn_graph* function from Monocle3 (Cao *et al*, 2019). **f)** Distribution of frags per cell recovered within bins across the T-cell activation trajectory. **g)** UMAP representation of only the activated T-cells colored by pseudotime bin (left) or stimulation type (right). **h)** Distribution of pseudotime positions of activated T-cells from either bead, or allogeneically (allo) stimulated conditions for each responder. **I**. Smoothed accessibility scores across pseudotime for sites found to vary over the trajectory (within activated T-cells). Top heatmap (Frac. Allo) displays the fraction of Activated T-cells within each pseudotime bin from MLR (Allo) conditions. **J**. Gene set analysis of genes with altered accessibility across pseudotime (within activated T-cells) using the Hallmark gene sets from MSigDB. Directionality scores for gene sets were determined using the *runGSA* function from the piano package(Subramanian *et al*, 2005).

After filtering for ATAC quality and hash enrichments, we resolved sample conditions for 18,567 filtered nuclei, spanning all experimental conditions and replicates (**Supp. Fig. 14c, Supp. Table S2**). Applying dimensionality reduction, we identified seven distinct clusters of chromatin states in our data (**Supp. Fig. 14b**). By integrating chromatin accessibility information with previously annotated single cell RNA-seq data from PBMCs (10X genomics, pbmc_10k_v3), we found nuclei within distinct clusters to be enriched for specific immune cell types (**Supp. Fig. 14c**). Combining the annotation enrichments for each cluster with accessibility scores for common immune markers enabled us to broadly annotate cell types within our experiment (**Fig. 4b, Supp. Fig. 14e**). Importantly, nuclei originating from “stimulator alone” conditions were disproportionately concentrated in clusters 3 and 6, which also showed the least enrichment for immune cell labels (**Supp. Fig. 14c**,**d**). Clusters 3 and 6 therefore likely represent irradiated PBMCs across MLR conditions and were excluded from downstream analyses.

Hash labels enabled us to define the proportion of immune cell-types recovered from each condition (**Fig. 4c**). By further subsetting cells by biological replicate, we further compared the proportion of each cell type recovered in stimulated conditions relative to unstimulated cultures (**Supp. Fig. 15a-d**). As expected, we observed an increase in the proportion of activated T-cells in the bead-stimulated condition and most allogeneic-stimulation conditions, but not autologously stimulated cells (**Fig. 4d**). Bead and allogeneic T-cell activation were further supported by flow cytometric analysis of cell proliferation for all conditions (**Supp. Fig. 16a-f**), illustrating that sciPlex-ATAC3 can detect meaningful cell type composition changes in heterogeneous specimens.

Given chromatin accessibility profiles for each cell, we strove to isolate changes to the chromatin of T-lymphocytes that are shared among donors, but enriched in allo-, as opposed to bead stimulation conditions. Upon TCR antigen recognition and co-stimulation, resting T-cells undergo dramatic chromatin remodeling, ultimately upregulating transcription of cytokines and cell cycle machinery(Rawlings *et al*, 2011). Using Monocle3, we identified a trajectory of chromatin changes between resting and activated T-cells (**Fig. 4e**). Consistent with general chromatin decondensation, we observed a large increase in recovered fragments per nucleus coinciding with progression towards the activated state (**Fig. 4f**). Surprisingly, when we examined activated T-cells alone, we noticed a more restricted localization of bead-activated T-cells, compared with those from allo-stimulated conditions (**Fig. 4g**), a pattern which holds across all individuals (**Fig. 4h**). To identify potential differences in gene regulation between stimulation types, we identified sites with altered accessibility across the trajectory within only activated T-cells. We found 3,973 sites which open or close across this axis (**Fig. 4i, Supp. Fig. 17a**), and opening sites were enriched for transcription factor binding motifs common to the AP-1 complex (**Supp. Fig. 17b**), a factor with established roles in T-cell activation(Rincón & Flavell, 1994). Similar TF enrichments were obtained when using sites identified through direct comparison of allo and bead-stimulated, activated T-cells (**Supp. Fig. 17d**,**e**). Furthermore, genes with increased accessibility scores across the activated T-cell trajectory were most associated with terms such as allograft rejection and IL2 signaling, in which T cells play a central role (**Fig. 4j**). Taken together, these results demonstrate that sciPlex-ATAC3 can distinguish between individual synthetic (bead-based) and allogeneically activated T cells based on their chromatin profiles alone, with genes and TFs canonically associated with T cell-mediated processes more accessible in allo-stimulated chromatin.

### Altered chromatin decondensation is a byproduct of chemically obstructed allogeneic T-cell activation

To investigate chemical immunosuppression of allogeneic T-cell activation, we next combined high-throughput chemical screening with cultured mixed lymphocyte reactions (**Fig. 5a**). Using donor D as the responder and donor A as an allogeneic stimulator, responder PBMCs were cultured separately with autologous, allogeneic or synthetic (bead) stimulators. Cells within each stimulation group were further cultured for five days with individual or combinations of immunosuppressive or epigenetic targeting compounds. Each treatment was performed at four distinct doses, including a vehicle control, and performed in biological replicate, totalling 192 conditions (**Supp. Fig. 18a**). Using sciPlex-ATAC3, all treatment wells were processed in a single batch, with 67,803 nuclei passing snATAC quality filters (**Supp. Fig. 18b**). 36,511 snATAC profiles remained after filtering for singlets and high confidence hashing. With the exception of dexamethasone, all chemical treatments reduced overall cell recovery with increasing dosage (**Supp. Fig. 18c**). Broad cell types were identified by integrating the vehicle treated cells from this experiment with annotated cells from our original MLR dataset (**Supp. Fig. 18d**) and were consistent with gene marker accessibility (**Supp. Fig. 18e**).

**Fig. 5:**
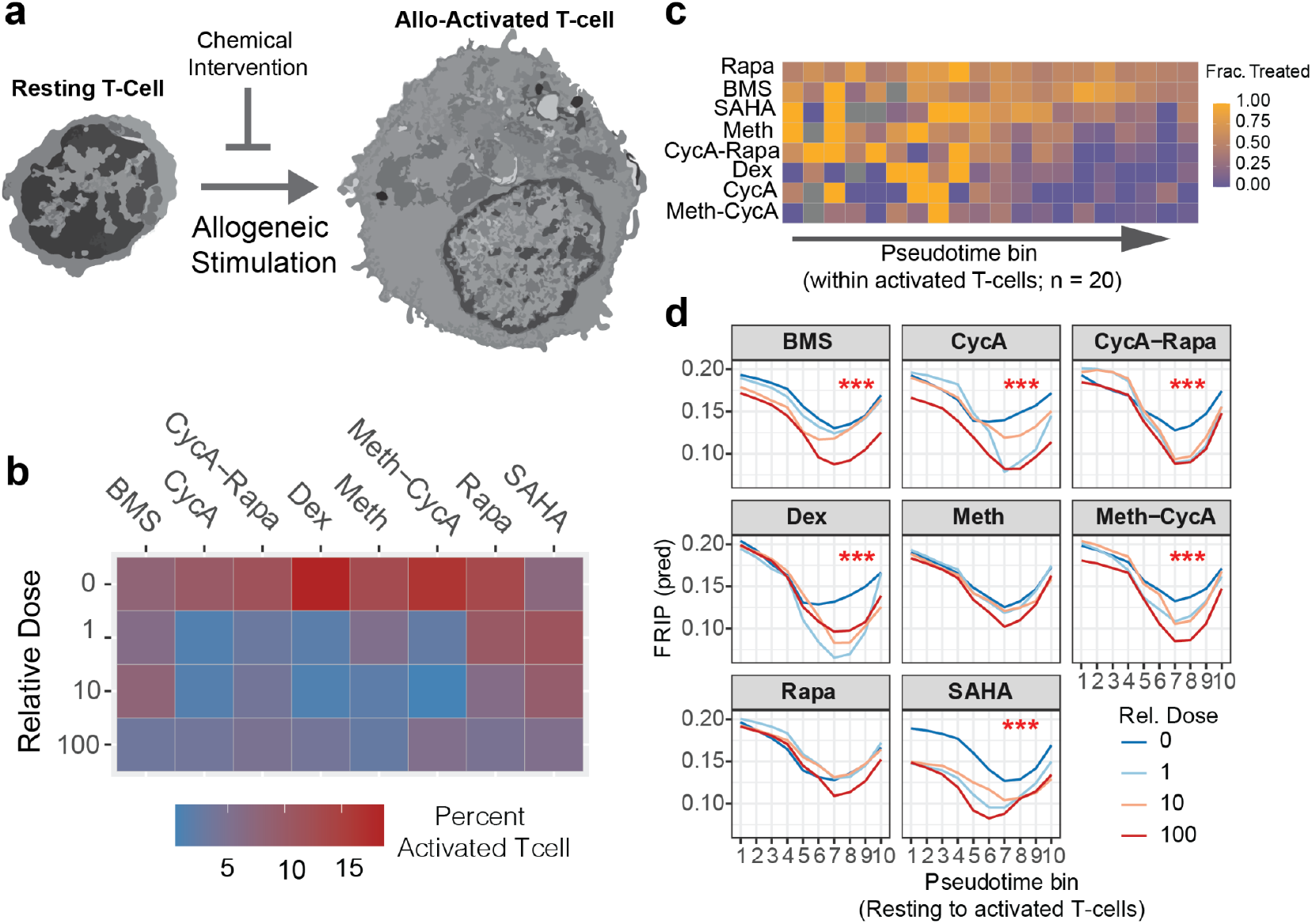
Altered chromatin decondensation is a byproduct of chemically obstructed allogeneic T-cell activation **a)** Illustration of using chemical intervention to block allogeneic activation of resting T-cells. **b)** heatmap showing the percent of activated T-cells (of all recovered cells) in allo-stimulated conditions treated with increasing dosage of eight different compounds/combinations. **c)** Heatmap showing the fraction of cells within each pseudotime bin treated with any non-vehicle dose of each compound. Rows/treatments were sorted by the sum of row values. **d)** Predicted FRIP scores for cells within each pseudotime bin from an interaction model of pseudotime position and drug dose. Red asterisks indicate a significant (*** = Pr < 0.0001) increase in goodness of fit for models including drug dose as an interacting term with pseudotime, compared with models using pseudotime alone (log ratio test).

Faceting chromatin profiles by stimulation type revealed only subtle impacts on the abundance of recovered chromatin states, while most chemical treatments severely altered the distribution of profiles, particularly at higher dosages (**Supp. Fig. 19a**). In particular, SAHA, BMS, and any treatment incorporating CycA increased the recovery of chromatin profiles enriched for dead/irradiated cells, consistent with toxicity of higher dosages (**Supp. Fig. 19b, 20**). Allo-stimulation significantly increased the recovery of activated T-cell profiles (**Supp. Fig. 21a**), which exhibited increased global chromatin accessibility compared with resting states (**Supp. Fig. 21b**,**c**).

To examine the impact of each chemical intervention on allogeneic T-cell activation, we focused our analysis on T-cells recovered from allo-stim conditions. Tellingly, increasing dosages of each treatment lessened the recovery activated T-cells in allo-responding conditions. Moreover, with clinically relevant compounds including dexamethasone, methotrexate and cyclosporin A, even the lowest doses prevented nearly all T-cell activation, when compared with auto stimulation (**Fig. 5b**). Very few of the allo-stimulated T-cell nuclei exposed to the explored compounds were found in the activated state (**Supp. Fig. 21d**). Nonetheless, by grouping allo-activated T-cells from each drug treatment by their activation trajectory positions, we identified variable degrees of inhibited progress towards the activated T-cell state among the drugs used. While combined treatment with methotrexate and cyclosporin A potently restricted the range of chromatin profiles, rapamycin, a compound commonly used to combat GVHD(Palmer *et al*, 2010) was much less impactful (**Fig. 5c**), even at the highest tested doses (**Supp. Fig. 21e**). Using the same trajectory-based groupings, we identified loci-specific chromatin changes along this allo-activated T-cell axis, with opening sites enriched for SMAD5 and KLF6 motifs regardless of timing (**Supp. Fig. 22a-f**). By inhibiting the majority of T-cell activation, drug treatments produced few obvious changes to specific chromatin-sites along the trajectory compared with untreated cells. However, the fraction of reads in peaks (FRIP) for nuclei along the T-cell activation trajectory were significantly influenced by most (6/8) compounds in a dose-dependent manner, pointing to increased accessibility within atypical regions as T-cells respond to allogeneic stimuli (**Fig. 5d**). Because nuclear decondensation in activated T-cells can proceed in the absence of key transcription factors(Lee *et al*, 2015), reduced FRIP values along the activation trajectory may point to deregulated chromatin remodeling downstream of these events in the rare instance of activation. Ultimately, sciPlex-ATAC3 reveals that diverse chemical regimens not only prevent the majority of resting T-cells from progressing towards allo-activated states, but also significantly disrupt canonical stimulation-dependent chromatin remodeling. These results likely reflect myriad disrupted cellular-processes for which additional phenotypic information will be invaluable.

## DISCUSSION

This work presents a straightforward, scalable strategy for hashing large numbers of samples for single-cell chromatin accessibility profiling. Notably, sciPlex-ATAC-seq is not without limitations. Extensive handling between barcoding reactions results in low overall nuclei yields (5-10%), and thus requires ample starting material. Moreover, omitting FACS-based sorting in sciPlex-ATAC3 to enhance nuclei recovery may lower data quality and increase sequencing costs. Several alternative approaches also enable multiplexed snATAC-seq, including dscATAC-seq (Mimitou *et al*, 2021; Lareau *et al*, 2019), ASAP-seq (Mimitou *et al*, 2021), and SNuBAR(Wang *et al*, 2021). While these approaches incorporate widely adopted droplet-based capture of nuclei, dscATAC-seq achieves scalability by incorporating an indexed-transposition reaction. ASAP-seq extends the concept of hashing cells with barcode-conjugated antibodies(Stoeckius *et al*, 2017), which can be co-assayed in droplets with accessible chromatin. Both dscATAC-seq and ASAP-seq attain impressive data quality, but either require many custom transposases, or expensive DNA-conjugated antibodies for sample multiplexing. Similar to our approach, SNuBAR uses unmodified DNA oligos to hash nuclei, but relies on their annealing to Tn5 sequences during transposition. By contrast, hashing for sciPlex-ATAC-seq is independent of transposition, enabling compatibility with single nuclei spatial profiling approaches(Srivatsan *et al*, 2021). SciPlex-ATAC-seq can be performed without expensive equipment, solely using commonly available reagents(Domcke *et al*, 2020). Moreover, aside from additional sequencing, increasing the scale of sciPlex-ATAC-seq experiments simply requires additional single stranded DNA oligos, making it attractive for HTS assays.

Applying sciPlex-ATAC-seq to a chemical screen, we observed drug and dose-specific effects of four compounds on the chromatin landscape of A549 cells, consistent with their mechanisms of action. Incorporating distal regulatory site information improved our ability to predict downstream effects of drugs on gene transcription, highlighting the mechanistic importance of non-coding regulatory elements(Luizon & Ahituv, 2015). By resolving chromatin profiles within individual cells we show that HDACis can induce heterogeneous chromatin responses, particularly at intermediate doses. Such variable responses to chemical inhibitors are not uncommon and can pose challenges in clinical settings(Shaffer *et al*, 2017). Uncovering the mechanistic basis and consequences of cell-to-cell variation in drug response will help guide the usage of therapeutically relevant compounds.

With the more scalable sciPlex-ATAC3, we also explored the immunogenic response of human PBMCs in mixed lymphocyte reactions. By performing and resolving multiple biological replicates for each condition, we were powered to detect altered T-cell proliferation and activation across MLR conditions. Bead-stimulated T-cells exhibited distinct chromatin profiles compared with activated T-cells from MLR conditions in a manner that was shared across donors. Interestingly, T-cell chromatin states specific to allo-activation appeared to reflect the activity of transcription factors such as AP-1 known to regulate genes involved in T-cell activation and differentiation(Rincón & Flavell, 1994), suggesting that bead stimulation only partially induces chromatin changes associated with T cell recognition of foreign cells. Upon activation, T-cells undergo a well characterized and rapid nuclear decompaction(Rawlings *et al*, 2011), a phenomenon also evident in our analyses(Rawlings *et al*, 2011; Gate *et al*, 2018; Calderon *et al*, 2019). Newly accessible regulatory elements within activated T-cells reflect the binding and activity of transcription factors which conspire to induce rapid proliferation, differentiation and signaling through altered gene expression programs(Gate *et al*, 2018; Calderon *et al*, 2019). By examining various chemical perturbations, we found that the majority of allo-T-cell activation can be potently inhibited via diverse pathways. However, for T-cells which managed to progress towards an activated-like state, most treatments also significantly disrupted global patterns of chromatin accessibility, possibly reflecting upstream mechanisms of action for these compounds. For example, treatment with SAHA may inhibit T-cell activation and proliferation by inducing acetate starvation and lowering intracellular pH (McBrian *et al*, 2013; Srivatsan *et al*, 2020). Calcineurin inhibition with cyclosporin A was previously found not to block large scale chromatin decompaction during T-cell activation(Lee *et al*, 2015), yet significantly altered the global distribution of accessible chromatin in our assay, perhaps as a symptom of reduced NFAT localization to the decondensed nucleus. Finally, unlike most examined conditions, the anti-inflammatory effects of rapamycin and methotrexate potently blocked allogeneic T-cell activation without detectably affecting global chromatin accessibility, emphasizing how drug action need not be directed through chromatin. Our analysis was limited to broad immune cell types, but resolving effects on T-lymphocyte subtypes will be crucial for understanding how allogeneic-stimulation and chemical immunosuppression might alter the fates of activated T-cells.

Ultimately, this work illustrates the potential value of sciPlex-ATAC-seq for dissecting how multicellular *in vitro* systems or disease models (e.g. organoids) respond to various perturbations. While continued development will be critical to address existing limitations and for cross-platform compatibility of sciPlex-ATAC-seq, the core technology is immediately applicable to a wide variety of biological applications aimed at exploring large parameter spaces. Moreover, as recently demonstrated (Chen *et al*, 2021; Fiskin *et al*, 2021; Ma *et al*, 2020; Mimitou *et al*, 2021; Cao *et al*, 2018), multimodal measurements provide unprecedented phenotypic information for individual cells and offer an attractive direction for future sciPlex-based assays.

## Acknowledgements

We thank Brian Beliveau and all members of the Trapnell, Shendure and Furlan labs for helpful discussions and feedback. In particular, we appreciate technical advice provided by Hannah Pliner and Silvia Domcke. **Funding:** Aspects of this work were supported by the NIH (UM1HG011586 and 1R01HG010632 to C.T. and J.S.; RC2DK114777 and R01HL118342 to CT; F32GM140502 to G.T.B.), American Cancer Society Mentored Scholar Award (to S.N.F), the Brotman Baty Institute for Precision Medicine, and the Paul G. Allen Frontiers Foundation (Allen Discovery Center grant to J.S. and C.T.). J.S. is an investigator of the Howard Hughes Medical Institute.

## Author Contributions

C.T. initiated and supervised the project. G.T.B., J.S., S.F and C.T. designed experiments. G.T.B., R.D., S.F., and R.G.G. performed experiments. G.T.B. analyzed the data. S.S. and J.L.M.F. provided initial guidance and technical support throughout. G.T.B., C.T., J.S. and S.F. wrote the manuscript with the support of all other authors.

## Data availability

Raw and processed data are available on GEO under accession number GSE178953. Any additional data will be made available upon reasonable request.

## Code availability

The scripts used for this manuscript are available on Github at https://github.com/gregtbooth/sciPlex-ATAC

## METHODS

### Hash Labeling Nuclei

Adherent cells grown in 96 well format were prepared by first aspirating the existing media. 50*µ*L of TrypLE (Termo-Fisher) was then added per well and the plate was incubated at 37°C for 15 minutes. After incubation, 150*µ*L of 1x DMEM (gibco) + 10% FBS(gibco) was added to quench the TrypLE reaction. The 200*µ*L cell suspension in each well was then transferred into a v-bottom 96 well plate, preserving the well orientations. Cells were then spun for 5 minutes at 300g to pellet cells before aspirating media. Cell pellets were washed with 100*µ*L 1xDPBS and then pelleted at 300g for 5 minutes. For suspension cells, well contents were first transferred to a v-bottom plate and then pelleted at 300g for 5 minutes. Cells were then washed in 200*µ*L 1xDPBS and spun down again before removing the DPBS. To isolate nuclei, pellets were then resuspended and gently pipetted up and down several times in 50*µ*L of either cold lysis buffer alone (10mM TrisHCl, 10mM NaCl, 3mM MgCl2, 0.1% Igepal, 0.1% Tween20, 1x Protease inhibitor (Thermo Pierce™ Protease Inhibitor Tablets, EDTA-free)), cold lysis buffer with 0.01% Digitonin (Promega), referred to as OMNI lysis buffer (Corces *et al*, 2017), or cold lysis buffer with 70*µ*M Pitstop 2 (Sigma) (Mulqueen *et al*, 2019). These conditions were compared in Supp. Fig. 1.

Single stranded DNA oligo labels (hashes) were then added to nuclei (aiming for approximately 1nMol hash molecules per 25,000 cells) in lysis buffer and incubated on ice for 5 minutes. Hashes have the sequence 5’-GTCTCGTGGGCTCGGAGATGTGTATAAGAGACAGXXXXXXXXXXBAAAAAAAAAAAAAAA AAAAAAAAAAAAAAA-3’, where ‘X’s represent the 10nt, well-specific hash ID. Ice-cold fixation buffer (1.5% Formaldehyde, 1.25x DPBS (Gibco)) was then added to samples to achieve a final formaldehyde concentration of 1% and mixed gently. Fixation was allowed to occur for 15 minutes on ice. At this point nuclei from different samples were combined and further steps performed on this single pool. Fixative was removed by spinning the pooled samples at 500g for 5 minutes. The pellet was then resuspended in nuclei suspension buffer (10mM TrisHCl, 10mM NaCl, 3mM MgCl2) + 0.1% tween20. Nuclei were pelleted again before being resuspended in freezing buffer (50 mM Tris pH 8.0, 25% glycerol, 5 mM Mg(OAc)2, 0.1 mM EDTA, 5mM DTT, 1x Protease inhibitor (Thermo Pierce™)) at a final concentration of 2.5 million nuclei/ml (two-level, sciPlex-ATAC), or 5 million nuclei/ml (three-level, sciPlex-ATAC3). Pooled samples were then flash frozen in liquid nitrogen and stored at -80°C.

### Co-capture of hash oligo and ATAC profiles with two-level sci-ATAC

Pooled, hash-labeled nuclei were thawed on ice, inspected for nuclei integrity, counted in the presence of Trypan blue (Gibco) and further adjusted to 2.5 million nuclei/ml if necessary. 2*µ*L nuclei were then distributed to all wells of a 96-well LoBind plate (Eppendorf). To capture hash molecules within each nucleus, 1*µ*L of 25*µ*M single-stranded DNA oligos (5’-TCGTCGGCAGCGTCAGATGTGTATAAGAGACAGNNNNNNNNXXXXXXXXXXTTTTTTTTT TTTTTTTTTTTTTTTTTTTTTVN-3’) were added to each well (‘X’s represent a well specific barcode while ‘N’s reflect the unique molecular index (UMI)). A table of the hash barcodes associated with samples for each experiment is provided in Supp. Table 3. The plate was then incubated at 55°C for 5 minutes and immediately returned to ice for 5 minutes. Capture oligos annealed to hash molecules were then extended by adding 3*µ*L of NEBNext High-Fidelity 2X PCR Master Mix to each well and incubating at 55°C for 10 minutes. Note that in Supp. Fig. 1 we also tested extension reactions with Superscript IV Reverse Transcriptase (Invitrogen) by adding 0.25*µ*L SSIV enzyme, 1*µ*L 5x SSIV reaction buffer, 0.25*µ*L water, and 0.25*µ*L 10mM DTT, and 0.25*µ*L 10mM dNTP to 3 uL nuclei. After extension 12*µ*L 2X tagmentation buffer (20mM Tris Ph 7.3, 10mM MgCl2, 20% DMF) and 4*µ*L of 4xCLB (40mM TrisHCl, 40mM NaCl, 12mM MgCl2, 0.4% NP40, 0.4% Tween20) was added to all wells. Note that for nuclei permeabilized with Pitstop2, the 4xCLB was supplemented with 280*µ*M Pitstop 2, while nuclei permeabilized and hashed in OMNI buffer received 4*µ*L of OMNI tag buffer (1.32x DPBS, 0.4% Tween20, 0.04% Digitonin). Finally, 1*µ*L of indexed Tn5 (Cusanovich *et al*, 2015; Amini *et al*, 2014) was added to each well and tagmentation was carried out at 55°C for 15 minutes before returning to ice. 96 uniquely indexed TN5-based reagents were prepared by mixing in equal parts all combinations of Tn5 complexes containing barcoded N5 ends (8x) and Tn5 complexes with barcoded N7 ends (12x). Unloaded Tn5 enzyme can be purchased from Diagenode (Cat:C01070010-10) and all sequences used for loading Tn5 are contained in Supp. Table 4. Tagmentation was stopped by adding 25*µ*L of ice cold 40mM EDTA + 1mM spermidine to all wells and then incubating at 37°C for 15 minutes. All wells were then pooled and DAPI was added to a final concentration of 3*µ*M for fluorescence-activated cell sorting (FACS). Using fluorescence based sorting, a limited number of cells (varied by experiment based on desired expected doublet rate) were distributed to each well of a new 96-well deep-bind plate containing 12*µ*L reverse cross-linking buffer (11*µ*L EB (qiagen), 0.5*µ*L 1%SDS, 0.5*µ*L 20 mg/mL Proteinase K (promega)) within each well. Cross-links were reversed by incubating plates at 65°C for 13.5 hours on a PCR block. PCR was then used to add a second round of well specific barcodes to both the hash labels as well as tagmented chromatin. To each well we added 3.65*µ*L Tween-20, 1.25*µ*L indexed Nextera P5 primer, 1.25*µ*L indexed Nextera P7 primer, and 18.125*µ*L NEBNext High-Fidelity 2X PCR Master Mix. PCR conditions were as follows:

72°C for 5 min
98°C for 30s

98°C for 10s
63°C for 30s
72°C for 1min
go to step 3 22X
72°C for 5 min
4°C

Amplified libraries from each well were then pooled and concentrated with Zymogen clean and concentrate kit (using 5X DNA binding buffer), before being eluted in 100*µ*L EB. To separate the hash library from the ATAC library, the concentrated, pooled library was run on a 1% agarose gel and gel purified. The hash library appears as a band of size 199 bp, while the ATAC library was cut from ∼200 - 3000bp. Gel extraction was performed with the Nucleospin PCR and Gel extraction kit and eluted in 50*µ*L (ATAC library), or 25*µ*L (hash library).

### Co-capture of hash and ATAC profiles with three-level sci-ATAC

Hash-labeled nuclei were thawed on ice, inspected for nuclei integrity, counted and further adjusted to 5 million nuclei/ml if necessary. 10*µ*L nuclei were then distributed to wells of a 96-well LoBind plate. To capture hash molecules within each nucleus, 2*µ*L of 25*µ*M of single-stranded DNA “capture” oligos (5’-TCGTCGGCAGCGTCAGATGTGTATAAGAGACAGNNNNNNNNTT+TTT+TTT+TTT+TTT+T TT+TTT+TTT+TTT+TTT+TVN-3’) were added to each well (‘N’s reflect the unique molecular index (UMI), ‘+T’ represents the presence of locked nucleic acids, which increase the melting temperature of the capture oligo annealed to hash oligo). For the experiment shown in Supp. Fig. 12, “enhanced” hash oligos had the following sequence: 5’-GTCTCGTGGGCTCGGAGATGTGTATAAGAGACAGXXXXXXXXXXCGGACGGTCGACATG GGATGAGAGGCCGCCGC-3. “Enhanced” capture oligos had the sequence: 5’-TCGTCGGCAGCGTCAGATGTGTATAAGAGACAGNNNNNNNNGC+GGC+GGC+CTC+TCA +TCC+CAT+GTC+GAC+CGT+CCG-3’ and were ordered with or without the presence of locked nucleic acids in positions indicated by “+”. The plate was then incubated at 55°C for 5 minutes and immediately returned to ice for 5 minutes. 35.5*µ*L of Tn5 reaction mix (25*µ*L 2X tagmentation buffer, 8.25*µ*L 1x DPBS, 0.5*µ*L 1%digitonin, 0.5*µ*L 10% tween-20, 1.25*µ*L water) was then added to each well. Finally, 2.5*µ*L TDE1 Tagment DNA Enzyme (Illumina) was added to each well (final volume = 50*µ*L). The plate was sealed with adhesive tape, and spun at 500g for 30 seconds. Tagmentation was then performed by incubating the plate at 55°C for 30 minutes. Tagmentation was stopped by adding 50*µ*L of ice cold 40mM EDTA + 1mM spermidine to all wells and then incubating at 37°C for 15 minutes. Using wide bore tips, all wells were pooled and tagmented nuclei were pelleted at 500g for 5 minutes at 4°C and the supernatant was removed. Nuclei were carefully resuspended in 500*µ*L 40mM TrisHCl, 40mM NaCl, 12mM MgCl2,+ 0.1% Tween-20 and spun again at 500g for 5 minutes at 4°C. Supernatant was aspirated and the pellet was resuspended in 110*µ*L 40mM TrisHCl, 40mM NaCl, 12mM MgCl2,+ 0.1% Tween-20.

5’ ends of tagmented chromatin and captured hash oligos within fixed nuclei were then phosphorylated via a polynucleotide kinase (PNK) mediated reaction. 110*µ*L of resuspended nuclei was mixed with 55*µ*L 10x T4PNK Buffer (NEB), 55*µ*L rATP (NEB), 110*µ*L nuclease-free water, 220*µ*L T4PNK (NEB), and 5*µ*L of the reaction mix was distributed to each well of a 96-well plate. The plate was then sealed, spun at 500g for 30 seconds, and then incubated at 37°C for 30 minutes.

Following kinase reactions, the first level of indexing was achieved by attaching indexed oligos specifically to the ‘N7-tagged’ side of tagmented chromatin and captured hash molecules. N7-specific ligations were performed by adding 10*µ*L 2X T7 ligase buffer, 0.18*µ*L 1000 *µ*M N7 splint oligo (5’-CACGAGACGACAAGT-3’), 1.12*µ*L nuclease-free water, 2.5*µ*L T7 DNA ligase (NEB), 1.2*µ*L 50*µ*M N7 oligo (5’-CAGCACGGCGAGACTNNNNNNNNNNGACTTGTC-3’, where ‘N’s represent a well specific index) directly to all wells containing the kinase reaction mixture (final well volume = 20*µ*L). The plate was then sealed, spun at 500g for 30 seconds, and ligation was carried out at 25°C for 1 hour. Ligations were stopped by adding 20*µ*L ice cold 40mM EDTA + 1mM spermidine to each well and incubating at 37°C for 15 minutes. Using wide bore tips, all wells were pooled into a 15ml conical tube and volume was increased by adding three volumes of 40mM TrisHCl, 40mM NaCl, 12mM MgCl2,+ 0.1% Tween-20. Nuclei were pelleted for 10 minutes at 500g and 4°C, and resuspended in 550*µ*L 40mM TrisHCl, 40mM NaCl, 12mM MgCl2,+ 0.1% Tween-20.

The second level of indexing was performed by ligating indexed oligos to the phosphorylated ‘N5-tagged’ side of tagmented chromatin and captured hash molecules. 5*µ*L of pooled, resuspended nuclei were thus distributed to all wells of a new 96-well plate. The second ligation reaction was then performed by adding 10*µ*L 2X T7 ligase buffer, 0.18*µ*L 1000*µ*M N5 splint oligo (5’-GCCGACGACTGATTA-3’), 1.12*µ*L nuclease-free water, 2.5*µ*L T7 DNA ligase, 1.2*µ*L 50 *µ*M N5 oligo (5’-CACCGCACGAGAGGTNNNNNNNNNNGTAATCAG-3’, where ‘N’s represent a well specific index) to all wells (final well volume = 20*µ*L). The plate was then sealed, spun at 500g for 30 seconds, and ligation was carried out at 25°C for 1 hour. Ligations were stopped by adding 20*µ*L ice cold 40mM EDTA + 1mM spermidine to each well and incubating at 37°C for 15 minutes. Using wide bore tips, all wells were pooled into a 15ml conical tube and volume was increased by adding three volumes of 40mM TrisHCl, 40mM NaCl, 12mM MgCl2,+ 0.1% Tween-20. Nuclei were pelleted for 10 minutes at 500g and 4°C, and gently resuspended in 500*µ*L EB buffer (Qiagen). For distribution to PCR wells, Nuclei were either stained with DAPI (3*µ*M final) and sorted into wells of a 96-well plate (185 nuclei/well) containing reverse cross-linking buffer (11*µ*L EB buffer (Qiagen) 0.5*µ*L Proteinase K (Roche), 0.5*µ*L 1% SDS), or counted and adjusted to a concentration of 1850/ml. 10*µ*L of diluted nuclei were distributed to all wells of a 96-well plate and 1*µ*L EB buffer (Qiagen) 0.5*µ*L Proteinase K (Qiagen), 0.5*µ*L 1% SDS was added to enable crosslink reversal. Plates were then sealed, spun at 500g for 30 seconds and crosslinks were removed by incubating plates at 65°C for 16 hours.

The third level of indexing is achieved through PCR. Therefore PCR mix containing 2.5*µ*L 25*µ*M P7 primer (5’-CAAGCAGAAGACGGCATACGAGATNNNNNNNNNNCAGCACGGCGAGACT-3’), 2.5*µ*L 25*µ*M P5 primer (5’-AATGATACGGCGACCACCGAGATCTACACNNNNNNNNNNCACCGCACGAGAGGT-3’), 25*µ*L NEBNext High-Fidelity 2X PCR Master Mix, 7*µ*L Water, 1*µ*L 20mg/mL BSA (NEB). Importantly, each well received a unique, well-specific, combination of P7 and P5 primers. PCR conditions were as follows:

72°C for 5 min
98°C for 30s

98°C for 10s
63°C for 30s
72°C for 1min
go to step 3 19X
72°C for 5 min
4°C

Amplified libraries from each well were then pooled and concentrated with Zymogen clean and concentrate kit (using 5X DNA binding buffer), before being eluted in 100*µ*L EB.

### 96-plex chemical treatments

From a single large culture, cells were washed with PBS and 25,000 A549 cells were seeded into each well of a 96-well flat-bottom culture dish (Thermo Fisher Scientific, cat no. 12-656-66). Cells were cultured in 100*µ*L DMEM (Gibco) containing 10% (v/v) FBS (Gibco) and 1% (v/v) Penicillin-Streptomycin (Gibco) for 24 hours prior to drug treatment. Drug dilutions were prepared at 100 times the desired dose such that all final treatments maintained a 0.1% concentration of vehicle in PBS. To initiate the treatments, 1*µ*L of the prepared compounds was added to each well to obtain the final target concentration. Cells from each well were harvested after 24 hours for SciPlex-ATAC-seq using the two-level approach with CLB lysis and tagmentation conditions as described above.

### Species mixing experiments

NIH-3T3 cells were purchased from ATCC (cat no. CRL-1658), while A549 were kindly provided by Dr. Robert Bradley (UW), and these two cell lines were used for all species mixing experiments. For each cell line, cells were separately cultured with DMEM (Gibco) containing 10% (v/v) FBS (Gibco) and 1% (v/v) Penicillin-Streptomycin (Gibco) in 10cm dishes to approximately 80% confluence. The adherent cells were washed with PBS, dislodged with TrypLE (Termo-Fisher) and diluted to desired concentrations in fresh culture medium. 50,000 cells in suspension were distributed to wells of a v-bottom 96-well plate (Thermo Fisher Scientific, cat no. 549935) for separate hashing of each cell line. For the pre- versus post-mixing experiment (Supp. Fig. 2), NIH-3T3 and A549 cells were resuspended and diluted in 1XDPBS, then either hashed and fixed separately (pre-mix), or mixed at equal proportions then hashed and fixed (post-mix) as described above.

### Mixed Lymphocyte Reactions and CFSE staining

Whole blood was extracted by venipuncture using 10ml Sodium Heparin tubes (BD Vacutainer) from four consenting volunteers (approved under IRB: FWA #00006878). PBMCs were then isolated from whole blood using Ficoll Paque Plus (GE Life Sciences Cat-17144002). Briefly, 15ml Ficoll was underlaid beneath 35ml diluted whole blood 1:1 with PBS in 50ml conical tubes. Tubes were centrifuged at 400g for 25 minutes without brakes for phase separation. The buffy coat containing PBMCs was then harvested into a new 50ml conical tube using a transfer pipette and resuspended with PBS to 50ml final volume. Harvested material was spun at 300g for 10 minutes and the supernatant was removed. Cells were resuspended in 3-5ml PBS for counting and transferred to 15ml conical tubes. PBMCs from each donor were further divided into responder and stimulator pools before being spun at 600g for 5 minutes and removing supernatant. Stimulator samples were resuspended at a final concentration of 4 × 10^6^ cells/mL in cTCM (RPMI (Gibco), 10% FCS (Gibco), 1X Pen-Strep (Gibco), 1X GlutaMAX (Gibco), 1.85*µ*L BME (Sigma) per 500ml of media) and irradiated (3500 cGy). Responders were resuspended at a concentration of 10 × 10^6^ cells/ml in PBS. All responder cells were labeled with carboxyfluorescein succinimidyl ester (CFSE) (ThermoFisher – Cat: C34554), regardless of experiment. Stock CFSE was added to responder cells to obtain a final concentration of 1*µ*M and gently vortexed. Responders were then incubated at 37°C in a water bath for 15 minutes with periodic gentle vortexing. CFSE labeling was then quenched by filling the 15 ml conical tube to the top with cTCM. Responder cells were then pelleted at 600g for 5 minutes. Cells were washed again with cTCM and spun as in the previous step. Responder cells were finally resuspended, counted, and adjusted to a final concentration of 4 × 10^6^ in cTCM before plating.

Responder cells from each of the four donors were plated in 96-well round bottom plates (Thermo) according to experimental layout in Supp. Fig. 13a, with each well receiving 400,000 cells in 100*µ*L. For bead stimulated conditions, only 80,000 responder cells were seeded per well (20*µ*L). An equal number of Irradiated stimulator cells were added to wells according to the experimental layout by adding 100uL (Supp. Fig. 13a). 100uL of extra cTCM was added to wells lacking stimulators to ensure all wells had 200*µ*L final volume. For bead stimulated conditions, MACSiBead particles with biotinylated antibodies against human CD2, CD3, and CD28 from the T Cell Activation/Expansion Kit, human (Miltenyi Biotech; Cat:130-091-441) were prepared according to manufacturer’s instructions. Beads were ultimately diluted such that each bead stimulated well received 8000 beads in a final volume of 200*µ*L cTCM. All conditions were harvested for sci-PlexATAC3 or fluorescent activated cell sorting-based measurements of CFSE staining five days after plating the experiment. Fluorescent cytometric analyses were performed using a BD FACSAria II machine.

### Chemical perturbations during mixed lymphocyte reactions

PBMCs were isolated from whole blood and divided into stimulator and responder pools as described above, except without CFSE staining. 2×10e5 responder cells were plated in each well of the allo and auto stim columns, while 1×10e5 responder cells were deposited into bead stimulation wells. Notably, unlike the multi-donor MLR experiment, for this experiment bead stimulations resulted in a lower recovery of activated T-cells compared to allo-activated conditions (Supp. Fig. 21a). This was a result likely due to expired reagents. For allo and auto stimulations, 2×10e5 irradiated stimulator cells were added to each well and all wells had a final volume of 200*µ*L with cTCM. Chemical treatments were initiated immediately after plating responders, stimulators and beads by adding 2*µ*L of the prepared compounds to each well to obtain the final target concentration (Supp. Table 5). Cells from each well were harvested for SciPlex-ATAC3 six days after treatments using the approach described above.

### Sequencing

For two-level sciPlex-ATAC experiments, which employ indexed Tn5 for the first level of chromatin barcoding, amplified ATAC and hash libraries needed to be sequenced separately. To sequence the two levels of barcodes introduced to chromatin via indexed-transposition PCR, a custom sequencing recipe was used as described previously (Cusanovich *et al*, 2015; Amini *et* al, 2014). Briefly, the following sequencing primers were used with Illumina Nextseq500 or Miseq platforms:

Read 1 (5’-GCGATCGAGGACGGCAGATGTGTATAAGAGACAG-3’)

Read 2 (5’-CACCGTCTCCGCCTCAGATGTGTATAAGAGACAG-3’)

Index 1(5’-CTGTCTCTTATACACATCTGAGGCGGAGACGGTG-3’)

Index 2(5’-CTGTCTCTTATACACATCTGCCGTCCTCGATCGC-3’).

The custom sequencing recipe uses “dark cycles” to prevent the 21-27nt constant region between two cell barcodes from crashing the sequencing software during index primed reactions. Therefore the following distribution of sequencing cycles was used: Read 1: 51 cycles, Read 2: 51 cycles, Index 1: 8 cycles+27 dark cycles+10 cycles, Index 2: 10 cycles+21 dark cycles+8 cycles. Index 1 captures the Tn5 barcode (T7), and then PCR barcode (P7). Index 2 captures the PCR barcode (P5), then Tn5 barcode (T5). The resulting indexed read is thus a 36nt sequence of the structure T7:P7:P5:T5.

The corresponding hash library was sequenced using standard primers and sequencing chemistry. Read 1 (18 cycles) captures the unique molecular identifier (8nt) as well as the PCR (p5) barcode (10nt). Read 2 (>=16 cycles) captures the hash ID. Index 1 (10 cycles) captures the PCR (p7) barcode and index 2 (10 cycles) captures the barcode from the in-well hash-capture oligo extension.

The custom sequencing recipe for sciATAC-seq3 has been described previously (Domcke *et al*, 2020) and was similarly followed here for 3-level sciPlex-ATAC (read 1: 51 cycles, read 2: 51 cycles, index 1: 10 cycles+15 dark cycles+10 cycles, index 2: 10 cycles+15 dark cycles+10 cycles). Because three level sciPlex-ATAC experiments result in hash molecules and chromatin fragments with identical barcodes and read structure the material was sequenced together for these experiments.

### Preprocessing sequencing data

The sequence processing strategy used here was established for sciATAC-seq2 and sciATAC-seq3, and has been described previously (Cusanovich *et al*, 2018; Domcke *et al*, 2020). Briefly, BCL files from Illumina sequencing were first converted to fastq files with bcl2fastq v2.16 (Illumina). Reads were filtered to include only those associated valid cell barcode combinations. Specifically, each of the four barcodes associated with each read pair, together comprising a cell ID, was required to be within an edit distance of 2 from expected barcode sequences. Valid barcodes were corrected for any errors and read pairs were trimmed with Trimmomatic-0.36 (Bolger *et al*, 2014) to remove adapter sequences. Trimmed reads were then aligned to either a hybrid human/mouse (GRCh38/mm9) genome (for species mixed samples) or the human (GRCh38) genome alone using bowtie2 (Langmead & Salzberg, 2012) with options “-X 2000 -3 1”. Properly aligned read pairs with a quality score of at least 10 were retained for downstream analysis using samtools with options “-f3 -F12 -q10”.

Because the hash label libraries from sciPlex-ATAC-seq2 must be sequenced separately (depicted in Fig. 1B), Preprocessing of these reads was performed separately from the corresponding chromatin libraries and follows a previously described approach (Srivatsan *et al*, 2020). Briefly, Illumina sequencing BCL files were converted to fastq files using bcl2fastq v.2.18 and reads with cell barcodes matching capture oligo indices within an edit distance of 2 bp were retained.

### Assigning sample labels based on hash reads

Either from separately sequenced hash libraries (sciPlex-ATAC-seq2), or the trimmed fastq files containing combined information (sciPlex-ATAC-seq3), bonafide hash labels were retrieved for sample assignment if the first 10nt of read 2 were an exact match to one of the hash indices employed for a given experiment. Hash reads were then grouped by their cell barcodes and collapsed based on UMIs within Read 1. Ultimately we obtain a vector of hash oligo UMI counts *h*_*i*_ for each nucleus *i* in the experiment.

Each nucleus was assigned to the treatment well from which it came by testing whether hash reads from each nucleus were enriched for a particular hash barcode. Hash UMIs within each nucleus were compared against a background distribution generated by averaging hash UMI counts within “nuclei” that did not exceed the minimum chromatin fragments per cell threshold (often attributed to nuclear debris). For each nucleus, hash UMI enrichment over background was assessed using a chi-squared test and labels were only assigned to nuclei with an adjusted (Benjamini-Hochberg) P-value < 0.05.

To use hash count information to distinguish singleton nuclei from multiplets or debris, enrichment scores were then calculated for each nucleus. Enrichment scores are defined as the ratio of the most abundant hash barcode to the second most abundant hash barcode within a given nucleus. Nuclei in which the most abundant hash was at least **α**-fold more abundant than any other hash barcode were considered singletons, while those not exceeding this threshold were considered multiplets or debris. **α** was determined on a per-experiment basis by examining the distribution of enrichment scores (see Fig. 1e) and selecting a value (at minimum an enrichment score of 2) distinguishing labeled and unlabeled cells. Ultimately, given the above criteria well-labeled nuclei met the following criteria: 1) At least 10 total hash UMIs with 2) an adjusted P-value < 0.05 enriched over background, and 3) a minimum enrichment score of **α**.

### sciPlex-ATAC-seq data processing and filtering

For each experiment, the final set of cells used for downstream analyses was required to meet the hashing criteria above, as well as pass quality thresholds regarding chromatin accessibility profiles. First, bam files containing aligned barcoded chromatin fragments were processed into Arrow files using the ArchR (version 0.9.5) package (Granja *et al*, 2021) with the following arguments; filterFrags = 500 and filterTSS = 3. Likely-doublets were then identified based on chromatin profiles using a modified version of the scrublet (Wolock *et al*, 2019) algorithm, which has been described in detail previously (Domcke *et al*, 2020). Briefly, doublets were simulated as the sums of random pairs of cells using the binarized cell-by-tile matrix (500bp genomic windows). LSI-based dimensionality reduction was performed on the log(TF)*log(IDF) transformed matrix. Note that the inverse document frequency (IDF) term in this transformation was derived prior to simulating doublets. Doublet scores reflect the fraction of simulated doublets within the set of nearest neighbors of a cell in the reduced dimension (dim = 49) space. Ultimately, the top 10% of cells from each experiment with the highest doublet scores were removed.

While the ArchR package includes a similar doublet calling algorithm (Granja *et al*, 2021), we were unable to obtain satisfying structure or clustering from the iterative-LSI dimensionality reduction framework on which ArchR is based. Therefore, we did not use ArchR for tasks requiring its implementation of dimensionality reduction.

### Processing and analysis of species-mixing experiments

To remove cells with low coverage, chromatin fragments per-cell cutoffs were selected based on the overall distribution within an experiment. Cells were also removed if they had fewer than 10 assigned hash UMIs. For each remaining cell barcode we tallied the number of reads uniquely aligning to human and mouse chromosomes (from the hybrid genome). Species collisions (likely doublets) were defined as cell barcodes with < 90% of fragments aligned to a single species. We then assessed how many species collisions were captured solely based on hash criteria defined above (i.e. low hash enrichment scores).

### Dimensionality reduction and trajectory analysis

Using the binarized cell by tile matrix (500bp genomic bins), UMAP projections of assayed cells were generated with Monocle3 to visualize the chromatin profiles in two dimensional space. Prior to dimensionality reduction, the cell by tile matrix was first filtered to only include genomic tiles accessible within at least 0.5% of assayed cells. This matrix was then preprocessed using the Monocle3 function ‘preprocess_cds’ with the following parameters, method = “LSI”, num_dimensions = 50. Dimensionality reduction and initial clustering was then performed using the ‘reduce_dimensions’ function with arguments: ‘reduction_method = UMAP’, ‘preprocess_method = LSI’, and ‘cluster_cells’ function with argument: ‘reduction_method = UMAP’ in monocle3. Note that for sciPlex-ATAC-seq3 experiments, prior to the dimensionality reduction step above, the first LSI component was removed, which improved cluster resolution as has been described previously for data of this type (Domcke *et al*, 2020).

### Defining accessible peaks

Using the initial cluster assignments defined through the monocle3 framework above, fixed-width (501bp) accessible genomic regions in each experiment were identified with ArchR. Specifically, replicate pseudobulk tracks were generated for Monocle3-defined clusters of cells using the ArchR function, “addGroupCoverages”. Peaks were then called using MACS2 (Zhang *et al*, 2008) from these simulated replicate coverage tracks using the ArchR function “addReproduciblePeakSet”, again grouping cells by Monocle3-defined clusters. Finally, using the resulting peak calls, a cell by peak matrix was generated using the ArchR function “addPeakMatrix”. Rather than calling peaks de-novo for the MLR perturbation screen dataset (Fig. 5), peaks called with the multi-donor MLR dataset (Fig. 4) were used, facilitating cross-experiment comparisons.

### Viability Curves

Viability curves shown in Supp. Fig. 4 were generated similarly to our previous work(Srivatsan *et* al, 2020). To model cell recovery as a function of drug dose, we first grouped per-well filtered single cell ATAC-seq cell counts by drug. These counts were then passed to the drm() function from the drc R package(Ritz *et al*, 2015) with the model formula ‘cells ∼ log_dose’ and the LL.4() model family function. This procedure fits a log-logistic model with the following form:

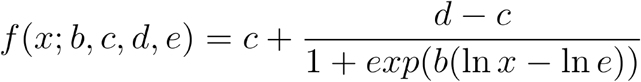

Above, the lower and upper asymptotic limits of the response are encoded by and, respectively. captures the steepness of the response curve and represents the half-maximal ‘effective dose’ (ED50).

### Dose response analysis

Using the peaks defined across cells from a given experiment, we applied a linear regression framework, implemented in Monocle3, to identify sites with accessibility altered in a drug and dose-specific manner. For each peak we fit the following logistic regression model to its accessibility across individual nuclei:

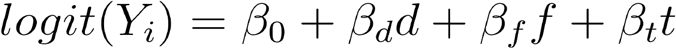

Where *Y*_*i*_ is a binary response variable for peak *i* (1 = “open” or 0 = “closed”),*d* is the log-transformed dose of the compound being evaluated, *f* is the log-transformed number of chromatin fragments within the nucleus, and *t* is the ArchR defined TSS enrichment score for the nucleus. For each model, we first subset cells to include only those relevant for determining a drug’s effect on a peak’s accessibility. To assess the effects on peak P in cells of type C when treated with drug D, we include in C, all cells treated with any dose of D. Additionally C includes cells treated with the vehicle control. For each drug D, peaks were only included in the analysis if accessible within at least 1% of C. The model above thus relates the accessibility of P across the subset of cells C. Peaks were determined to be differentially accessible if their fitted models include a dose term β_*d*_ that is significantly different from zero as assessed by a Wald test (Benjamini-Hochberg adjusted). Prior to correcting for multiple testing, values for terms are pooled across all compounds and all analyzed peaks prior to correction for multiple testing.

### Pseudodose trajectories

Pseudodose trajectories for each drug were generated by repeating the dimensionality reduction procedure described above on the peak-by-cell matrix restricted to the subset of cells within each drug-treatment group including vehicle controls. After performing dimensionality reduction and running UMAP, we fit a principal graph to the data using the learn_graph() function in Monocle3. We defined the origin (roots) of the trajectory by first assigning each cell to their nearest graph node. Nodes for which a majority of the assigned cells were treated with vehicle were called root nodes. For all remaining cells, their pseudodose ***Ψ*** was defined as the geodesic distance between their assigned node and a root node. For comparisons between pseudodose values and actual treatment doses, cells were grouped based on their pseudodose values using k-means clustering (k = 10) and the mean value was assigned to all cells within the group.

As described previously(Pliner *et al*, 2018), to generate smoothed accessibility profiles for each peak (as in Fig. 3D), grouped cells were further subdivided into groups containing at least 50 and no more than 100 cells. Aggregate accessibility profiles were then generated for each group from binary data of individual cells, creating a matrix A such that A _ij_ contains the number of cells in group j for which peak i is accessible. Importantly, we preserved the average group pseudodose value ψ_j_ and overall cell-wise accessibility S_j_ for cells in each group i. We then fit the following model to the binned accessibility within each group:

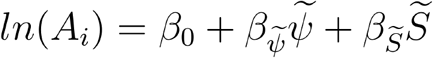

Where A_i_ is the mean of a negative binomial valued random variable of cells in which site i is accessible. The 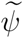 and 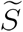 variables were smoothed with a natural spline during fitting. Smoothed values were retrieved for each peak and pseudodose bin via the model_predictions() function in Monocle3.

### Motif enrichment analysis

A binary peak by motif matrix was generated for our experiments using the ArchR function *addMotifAnnotations* with the argument *motifSet = cisbp*. Using this matrix, we then applied a logistic regression model that uses the presence or absence of individual motifs in each peak to predict whether the site had a particular accessibility trend (opening, closing, or unchanged). The model had the formula:

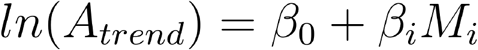

Where *ln*(*A*_*trend*_)is a binary indicator of a site’s accessibility trend (i.e. opening or not) and represents the presence or absence of motif *i* within a site. To find motifs enriched within opening, closing and unchanged sites, this model was applied separately for each accessibility trend identified within each treatment condition. A motif *i* was identified as enriched within peaks of a given trend if its *β*_*i*_*M*_*i*_ was significantly predictive of a site’s accessibility trend, as determined by two-tailed z-test (Benjamini-Hochberg adjusted p <0.05).

### Integration of single cell ATAC-seq and single cell RNA-seq data

To directly relate single cell chromatin accessibility profiles with previously determined transcriptomes from the same chemical screen(Srivatsan *et al*, 2020), we used the ArchR function *addGeneIntegrationMatrix* which employs a label transfer algorithm developed previously(Stuart *et al*, 2019). Using the gene score matrix calculated by ArchR and the previously generated RNA count matrix for the chemical screen, we performed an unconstrained integration (i.e. allowing cells to find closest match to any cell in the RNA data) and assigned predicted treatment labels based on that of the resulting assigned cell transcriptome. To directly compare drug effects on gene activity (as measured by ATAC) and gene transcription (as measured by RNA), we fit a genes (library size-factor adjusted) gene score (ATAC) or UMI count (RNA) recorded from each nucleus with a generalized linear model:

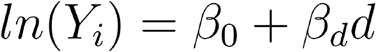

Where *Y* is a quasipoisson-valued random variable and *d* is the log-transformed dose of the compound being examined. We applied this model to our data (either the gene score matrix or RNA count matrix) as described above for the dose response analysis of peak accessibility. Ultimately, we compared the coefficients for d attained for each gene when we used the gene score matrix versus the RNA count matrix.

### Identifying cicero connections

We identified co-accessible sites using the Cicero R package for Monocle3 (version 1.3.4.5)(Pliner *et al*, 2018). For the chemical screen experiment cicero was run using all filtered cells from the experiment. To calculate co-accessibility scores, Cicero cell data set objects were generated using the *make_cicero_cds* function with default parameters on the reduced dimension cell by peak matrix and corresponding UMAP coordinates determined by Monocle3 as described above. Co-accessibilty scores were then calculated using the *run_cicero* function with default parameters. When calculating promoter versus distal pairs, any site within 500bp of an annotated TSS was labeled as a promoter, while all other sites were labeled as distal.

### Regression models for gene expression

Similar to the approach used previously(Pliner *et al*, 2018), we fit two regression models that predict, for each promoter region, the coefficient β_*d*_*d*(defined above) describing the effect of drug dose on gene expression (measured by sciPlex-RNA-seq). We excluded promoters which did not have an accessible promoter (defined as having an ATAC peak within 500 bp of the TSS) or did not have a drug-dose coefficient significantly different from 0. For each drug, the model was applied to the following number of promoters: SAHA: n= 4101, Dex: n= 1456, Nutlin3A: n = 1280, BMS: n = 1181. For the first set of models (“promoter motifs”) the features are binary values for each motif, indicating whether it is present within at least one accessible peak overlapping the promoter (TSS +/- 500bp). The second set of models (“promoter and distal motifs”), the features are the promoter motif indicators plus a second set of real-valued variables encoding information regarding distal sequence motifs. Specifically, the distal motif variable for each motif and TSS equals the highest cicero co-accessibility score from a promoter peak for that TSS to any connected distal peak which possesses that motif. If no connected distal peak contains the motif, the distal motif variable is assigned a value of 0. The models were trained using elastic net regression. Peaks were required to have a co-accessibility score of 0.1 or greater to be considered connected.

### Modeling fraction of reads in peaks (FRIP) over drug-treated T-cell activation trajectory

FRIP values were determined for each cell as the total number of fragments overlapping called peaks divided by the total number of fragments recovered. To test whether the drug dose variable significantly contributed to altered FRIP values along the T-cell activation trajectory, we fit the following model to the FRIP values for each cell within a drug treatment group:

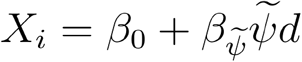

Where *X*_*i*_ is the FRIP score for nucleus i, and assumed to be gaussian distributed. The variable 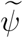 represents a cell’s pseudotime trajectory position and was smoothed with a natural spline during fitting. The variable *d* represents drug dose. We then applied the log ratio test (LRT) to evaluate whether the above model was a better fit than the following reduced model, which excludes the dose interaction term d:

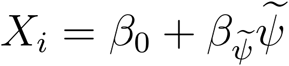

FRIP was determined to be significantly affected by treatment dose if the full model was found to significantly improve prediction by LRT ((pr > Chisq) < 0.05). Separate models were fit and examined for cells from each drug treatment group and their corresponding vehicle-treated cells.

