## Supplemental Figures and Legends for "High-Capacity Sample Multiplexing for Single Cell Chromatin Accessibility Profiling"

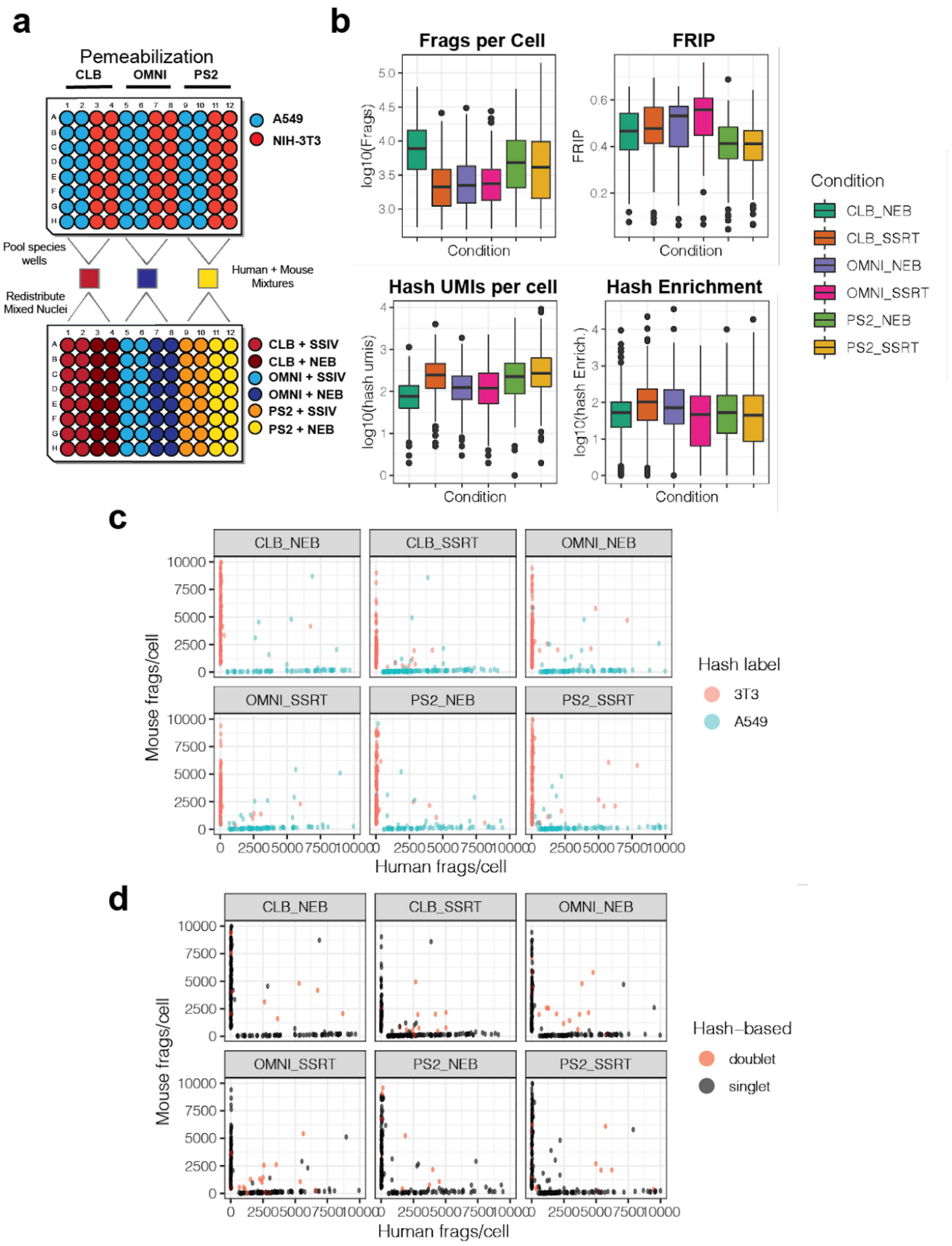

a
Pemeabilization
CLB
OMNI
PS2
A549
NIH-3T3
Pool species wells
Human + Mouse Mixtures
Redistribute Mixed Nuclei
CLB + SSIV
CLB + NEB
OMNI + SSIV
OMNI + NEB
PS2 + SSIV
PS2 + NEB
b
Frags per Cell
FRIP
Hash UMIs per cell
Hash Enrichment
log10(Frags)
FRIP
log10(hash umis)
log10(hash Enrich.)
Condition
Condition
CLB_NEB
CLB_SSRT
OMNI_NEB
OMNI_SSRT
PS2_NEB
PS2_SSRT
c
Hash label
3T3
A549
Mouse frags/cell
Human frags/cell
d
Hash–based
doublet
singlet
Mouse frags/cell
Human frags/cell

**Supplementary Figure Fig. 1:** Comparison of various nuclei permeabilization and hash extension strategies. **a)** Schematic of experimental design for comparing 6 distinct library preparation procedures in parallel. Human (A549) and mouse (NIH-3T3) cells were distributed into a 96-well culture plate (Top) where nuclei were isolated and permeabilized with one of three solutions. After permeabilization nuclei in each well were incubated with unique hash labels and then fixed. After fixation nuclei were pooled based on their permeabilization buffers, resulting in 3 pools of hashed nuclei from both species. Species-mixed nuclei were then distributed into a second 96-well plate, as shown, where hash extension was performed with one of two enzymes/reagents. **b)** Boxplots depicting distributions of quality metrics for cells exposed to distinct library preparation strategies. **c)** Scatter plots of unique human and mouse fragments per cell, faceted by library preparation conditions. Colors show species assignments based on the dominating associated hash label for each cell. **d)** Scatter plots of unique human and mouse fragments per cell, faceted by library preparation conditions. Colors identify doublets assigned based on the inability to confidently assign a dominant associated hash label within a cell.

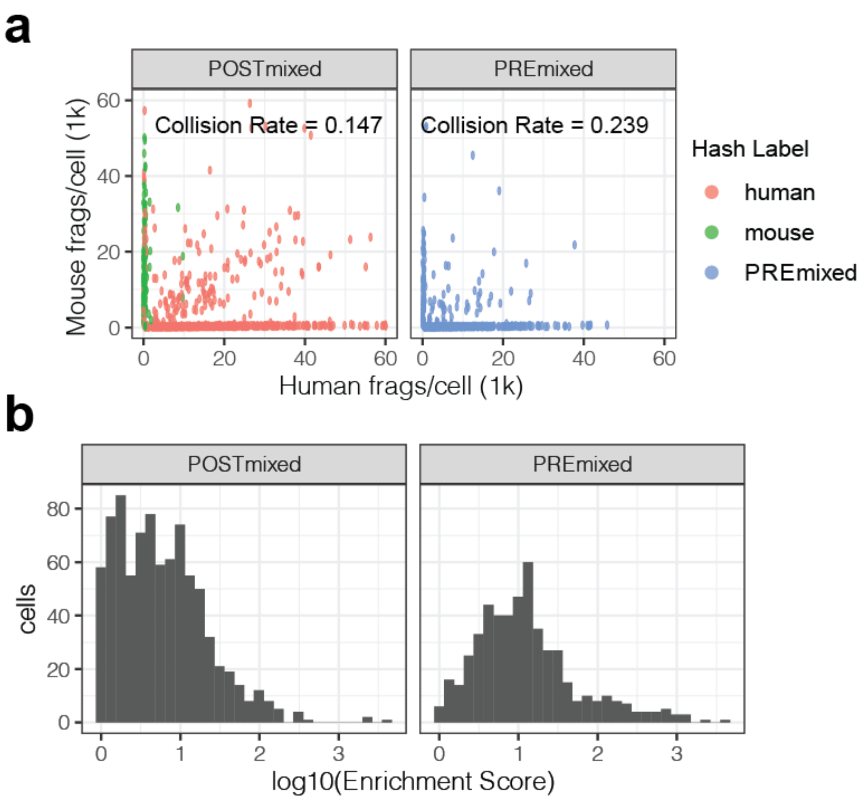


**Supplementary Figure Fig. 2:** Fixation does not increase doublet formation. **a)** Scatterplots of unique mouse and human fragments recovered per cell from samples where cells were mixed before (Pre) or after (Post) hashing and fixation. **b)** Hash enrichment scores from pre and post-hashing pooled cell-types.

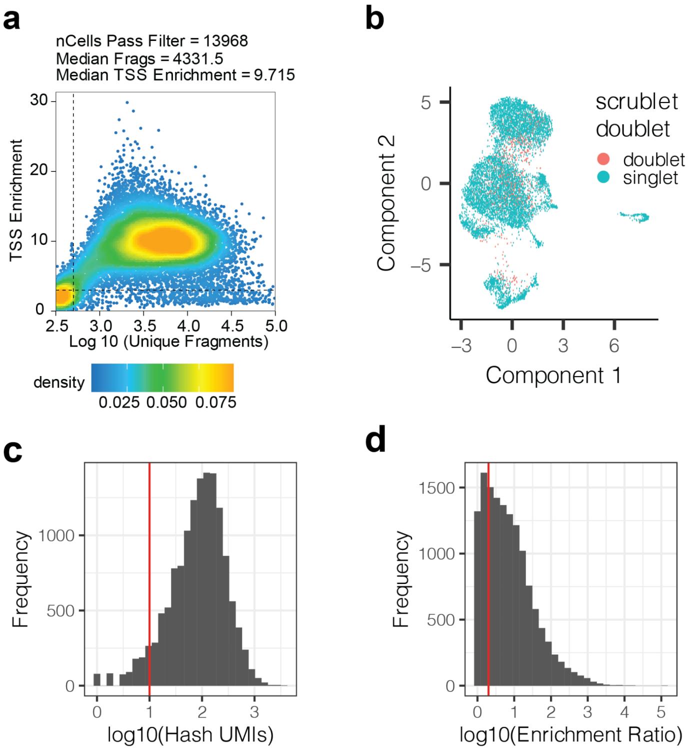


**Supplementary Figure Fig. 3:** Hashing is compatible with high throughput chemical epigenomic screens. **a)** Scatter plot showing the relationship between recovered fragments per cell and TSS enrichment. Dotted lines represent baseline per-cell cutoffs for each value. Cells passing these cutoffs were further filtered based on hashing (see methods) **b)** UMAP position of remaining doublets, after hash-based filtering, identified with a modified version of scrublet (Wolock *et al*, 2019) for scATAC data (Domcke *et al*, 2020). **c)** Distribution of hash umi counts recovered per cell. Vertical line represents a filter cutoff (< 10) to remove low quality cells. **d)** Distribution of hash enrichment scores for all cells. The hash enrichment score is defined as the number of hash umis recovered from the most abundant ID divided by the second most abundant ID within a cell. Vertical line represents a filter cutoff (enrichment > 2) to remove low quality cells.

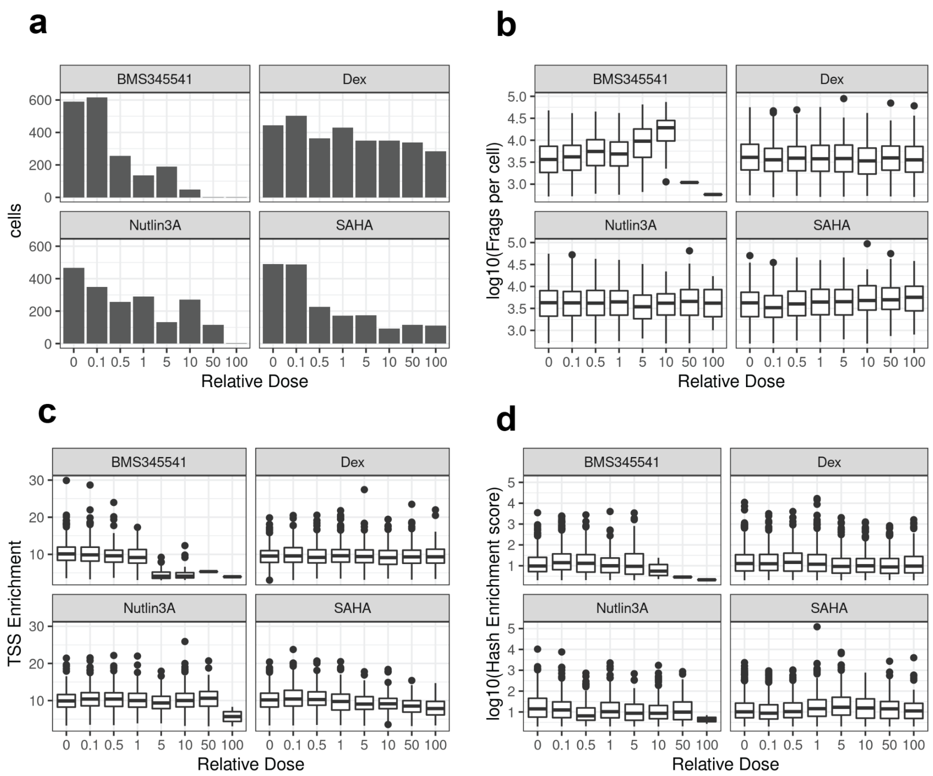


**Supplementary Figure Fig. 4:** SciPlex-ATAC-seq metrics by chemical treatment. **a)** Number of cells (passing filters) recovered per condition. **b)** Distributions of chromatin fragments recovered per cell by condition. **c)** Distribution of TSS enrichment values per cell by condition. Significance of relationships between drug dose and TSS enrichment was evaluated using the 'glm' package in R with the following model: 'TSSenrichment ~ ln(dose) + frags_per_cell' (Dex. dose coefficient = -0.02, p = 0.11; Nutlin3A dose coefficient = 0.019, p = 0.388). **d)** Distribution of hash enrichment scores per cell by condition. The hash enrichment score is defined as the number of hash umis recovered from the most abundant ID divided by the second most abundant ID within a cell. Vertical line represents a filter cutoff (enrichment > 2) to remove low quality cells

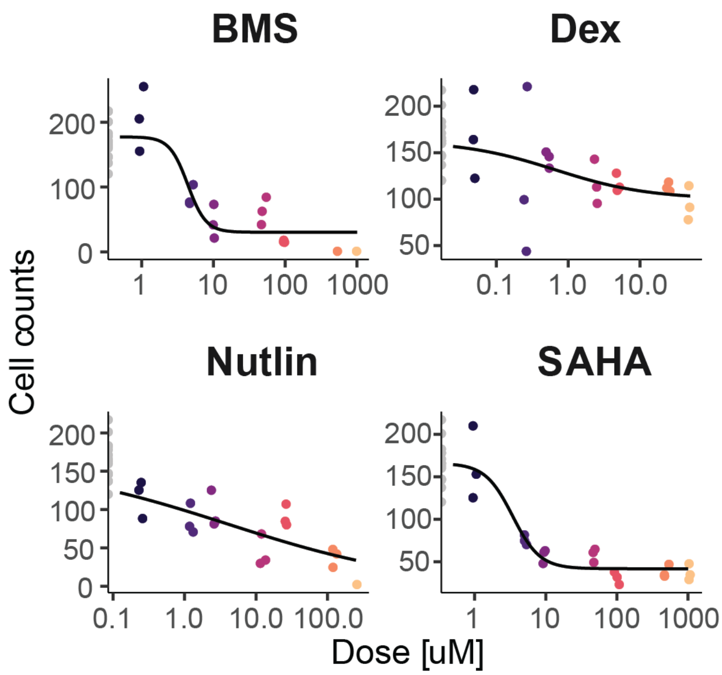


**Supplementary Figure Fig. 5:** Viability curves can be fit to the number of cells recovered at each treatment dose. Curves were fit to cell counts recovered from each dose treatment of BMS345541, dexamethasone, nutlin-3a and SAHA as previously described(Srivatsan *et al*, 2020).

| Compound | Distal_closing | Distal_opening | Promoter_closing | Promoter_opening |
|---|---|---|---|---|
| BMS | 66 | 117 | 1 | 20 |
| Dex | 103 | 666 | 7 | 16 |
| Nutlin | 2 | 62 | 0 | 7 |
| SAHA | 956 | 822 | 186 | 68 |

**Supplementary Table 1:** Number of differentially accessible sites, either promoter proximal or distal, found to open or close in response to each treatment. The distal category includes all non-promoter element types (distal, intronic, exonic). Opening and closing corresponds to sites with regression coefficients for drug dose > 0, or < 0, respectively.

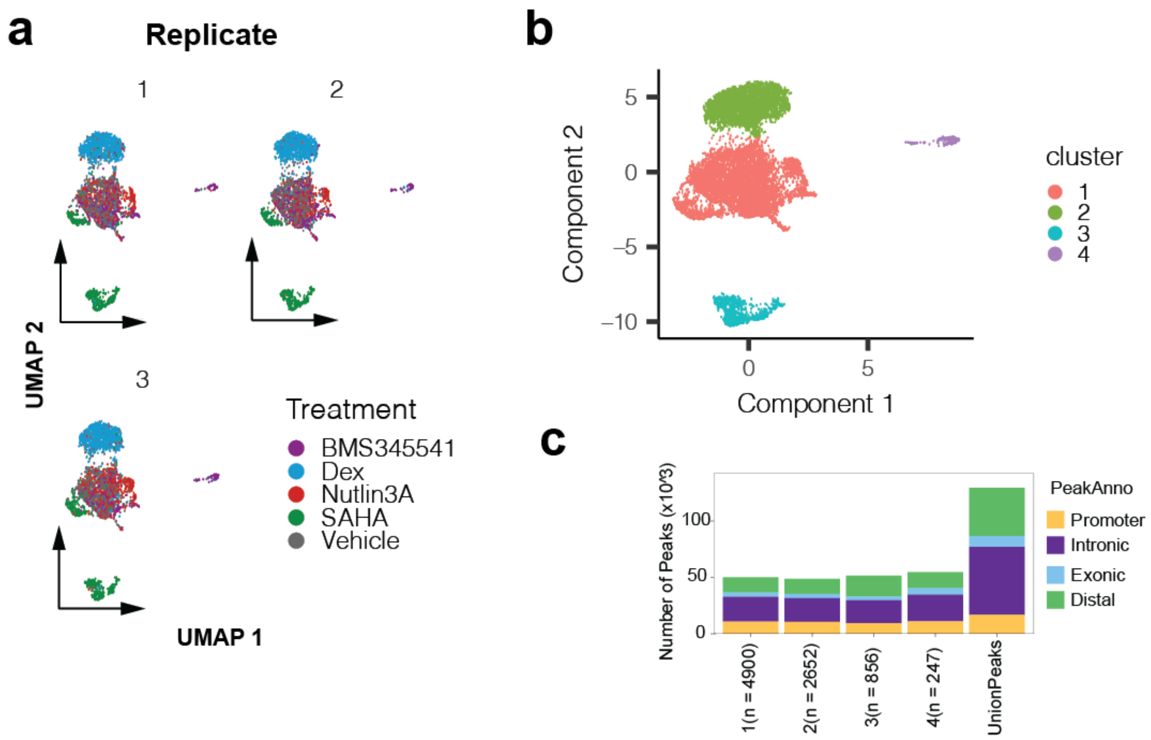


**Supplementary Figure Fig. 6:** Reproducible clustering facilitates accessible peak calling. **a)** UMAP projections of chromatin profiles from chemical screen, faceted by replicate wells for each treatment. **b)** UMAP projection of chromatin profiles colored by clusters identified with Monocle3 (Cao *et al*, 2019). **c)** Number of accessible peaks identified from each cluster with ArchR (Granja *et al*, 2021) and their annotations (UCSC).

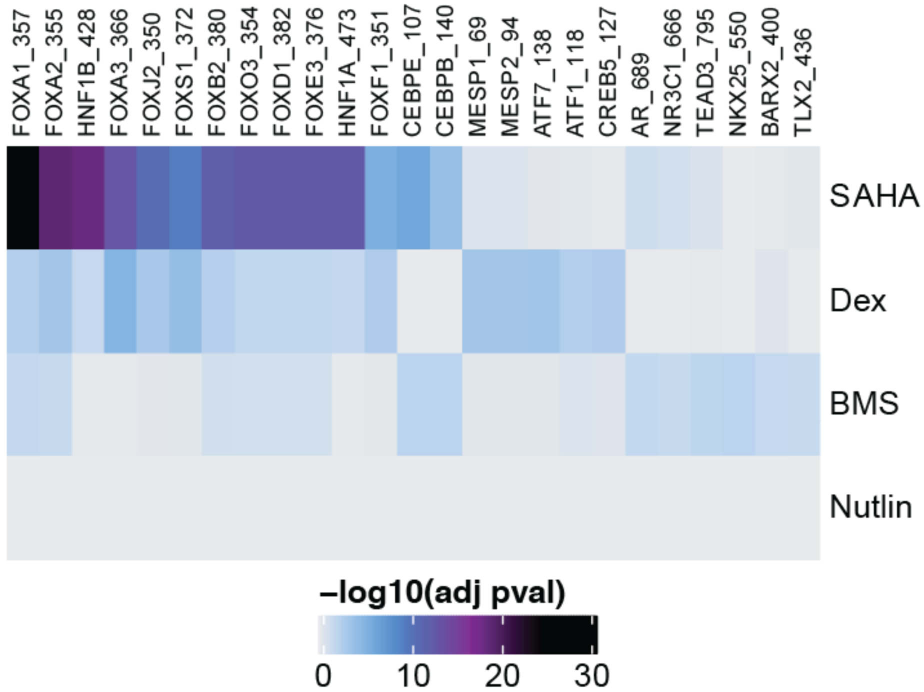


**Supplementary Figure Fig. 7:** Drug-specific motif enrichments are consistent with known compound mechanisms of action. Heatmaps depicting up to 10 most significantly enriched motifs per drug within peaks found to close with treatment.

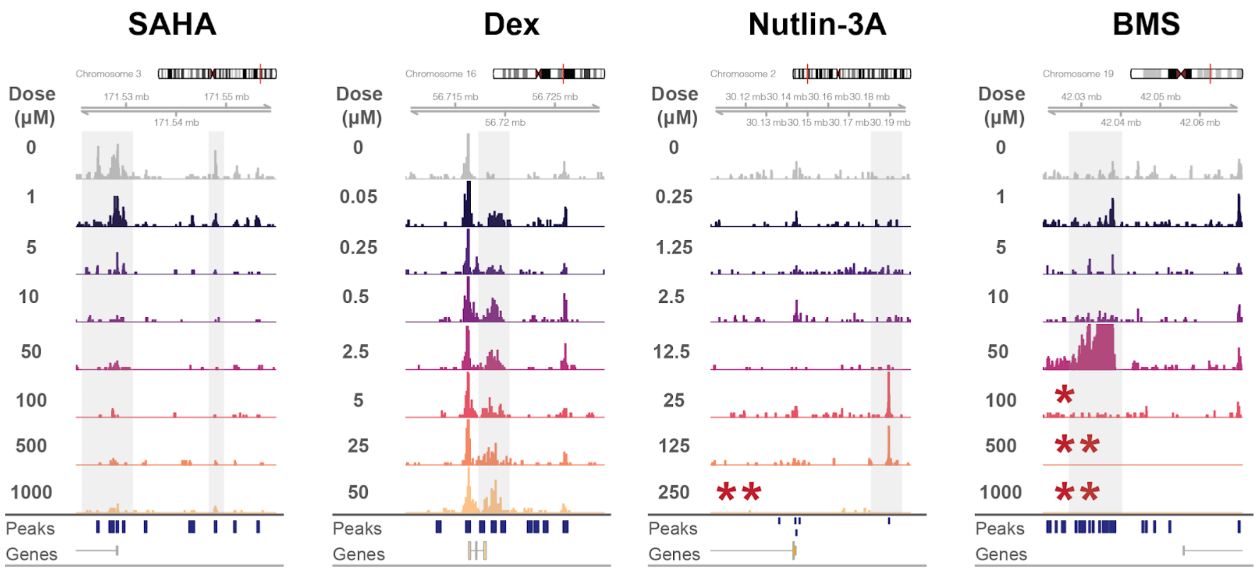


**Supplementary Figure Fig. 8:** Browser tracks of pseudobulk accessibility read coverage for cells grouped by treatment dose for each compound used. Y-axes are the same for all tracks (range 0-20) and represent pseudo bulk read coverage, normalized by reads within promoters for each group. Regions represent loci identified as significantly altered for individual drugs. Asterisks indicate few cells were contained within the group ( * < 50, ** < 10).

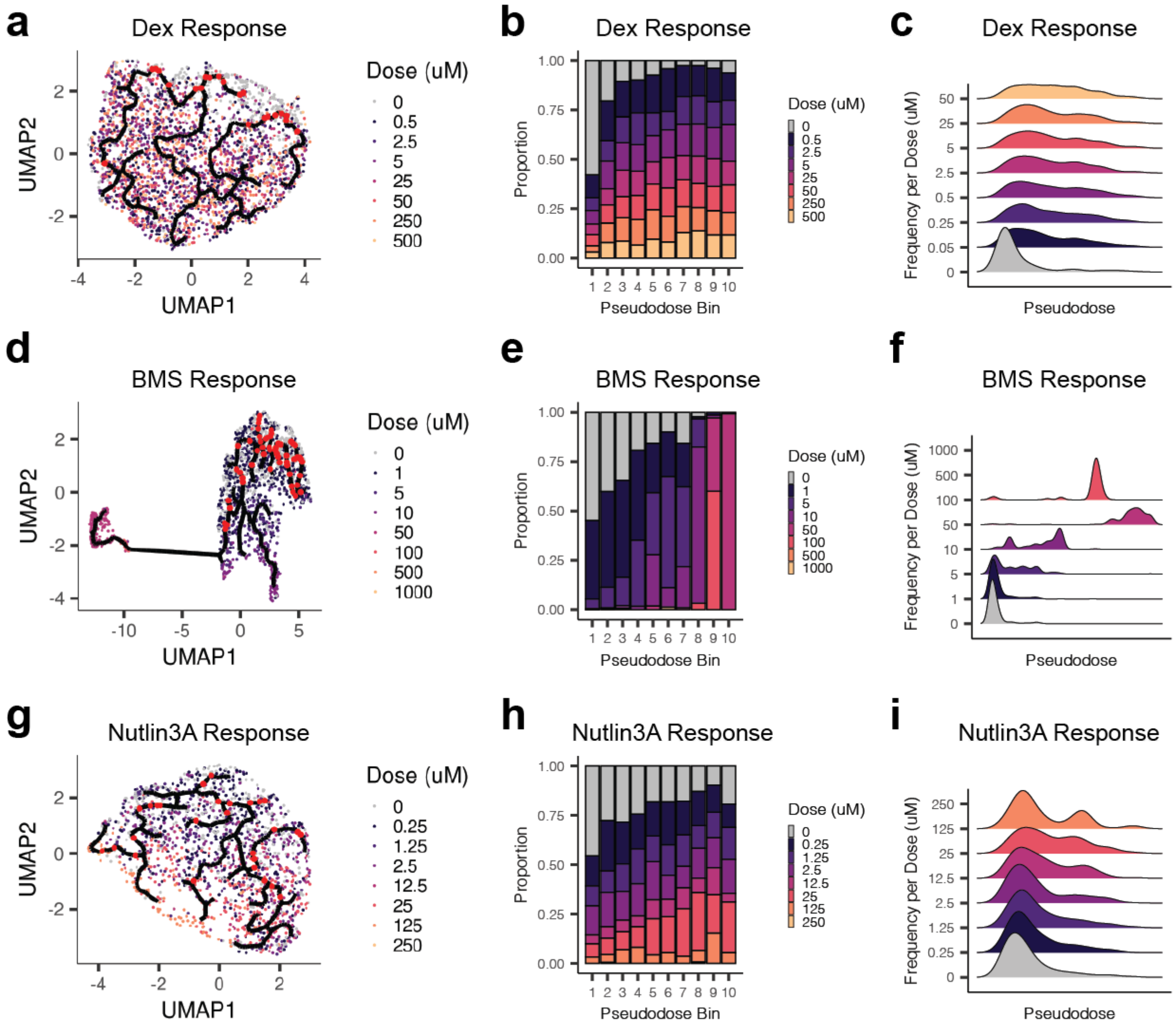


**Supplementary Figure Fig. 9:** Trajectory analysis reveals progression of chromatin state changes in response to drug treatment. **a)** UMAP embedding of cells from the dexamethasone treatment group (including vehicle controls), colored by dose of treatment. Red dots reflect positions identified as root nodes for the drug response trajectory (see methods). **b)** Barplots depicting the proportion of cells treated with each dose of dexamethasone within each pseudodose bin. **c)** Distribution of cells treated with each dose of dexamethasone across pseudodose chromatin states. **d-f)** Same as A-C but for cells treated with BMS345541. **g-i)** Same as A-C but for cells treated with Nutlin-3A.

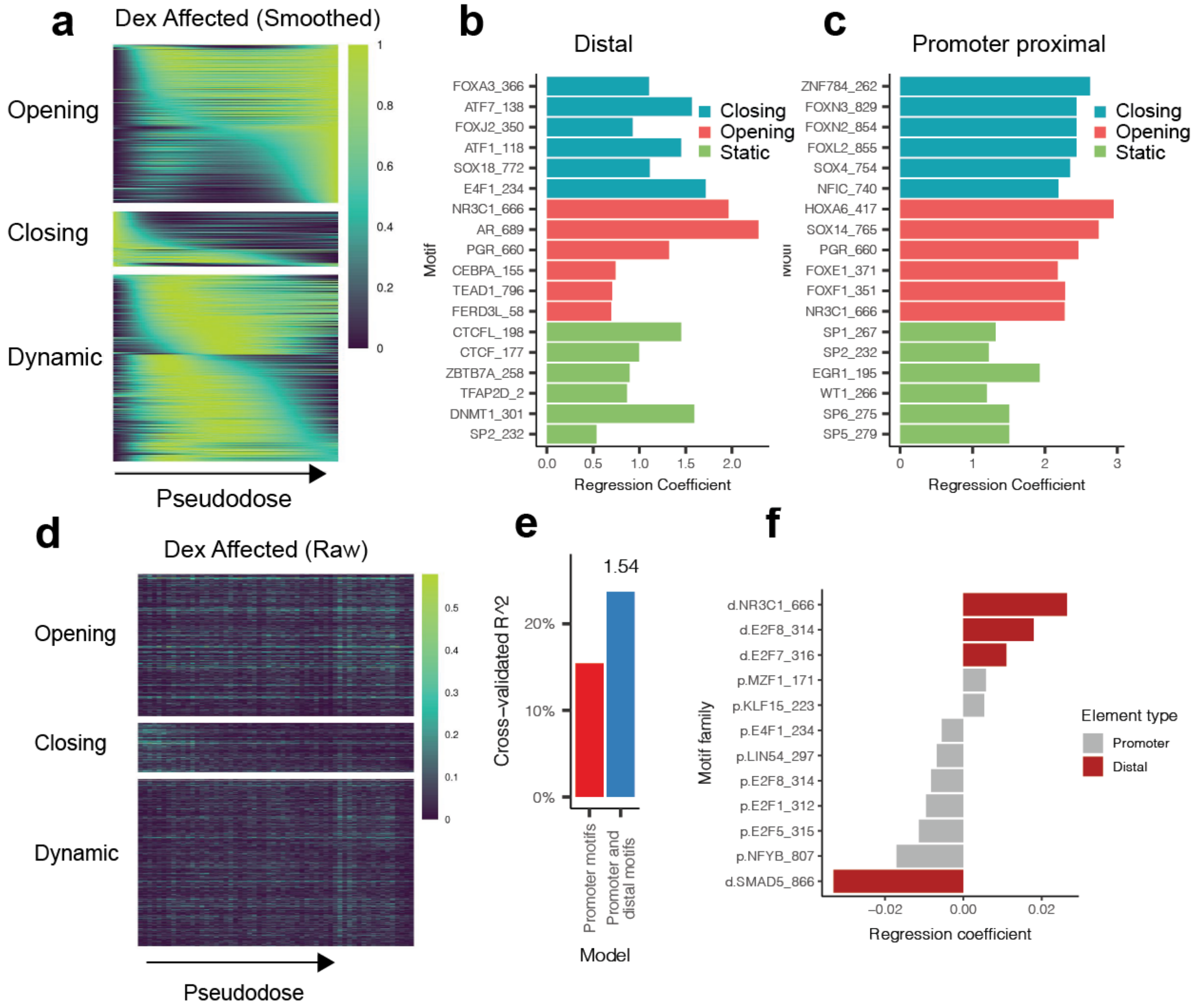


**Supplementary Figure Fig. 10:** Dex-responsive chromatin states predict transcriptional response. **a)** Smoothed accessibility scores across Dex-pseudodose for three classes of identified differentially accessible sites (opening, closing, and dynamic). Closing and opening sites were defined as sites with a maximum accessibility score occurring within the first or last 20 pseudotime bins (out of 100 bins), respectively. Dynamic sites have a maximum score in the intervening pseudotime bins. **b)** Top motifs explaining whether a distal Dex-DA site is classified as closing (blue), opening (red) or static (non-DA, green). **c)** same as E, but for Dex-DA sites overlapping gene promoters. **d)** Raw accessibility scores (fraction cells accessible in each bin) across Dex-pseudodose for three classes of identified differentially accessible sites (opening, closing, and dynamic). **e)** Dex-responsive DE Gene expression variation explained by models taking into account sequence elements within promoters alone, or within promoters and distal co-accessible sites. The value (1.54) above the right bar reflects the fold increase in predictive power with models which include distal co-accessible sites. **f)** Sequence elements with largest regression coefficients from promoter (grey) or distally connected sites (red) when predicting Dex-affected gene responses.

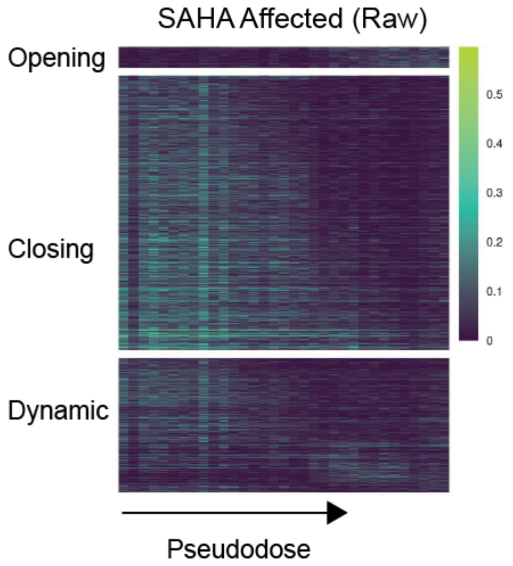


**Supplementary Figure Fig. 11:** Raw accessibility scores (fraction cells accessible in each bin) across SAHA-pseudodose for three classes of identified differentially accessible sites (opening, closing, and dynamic)

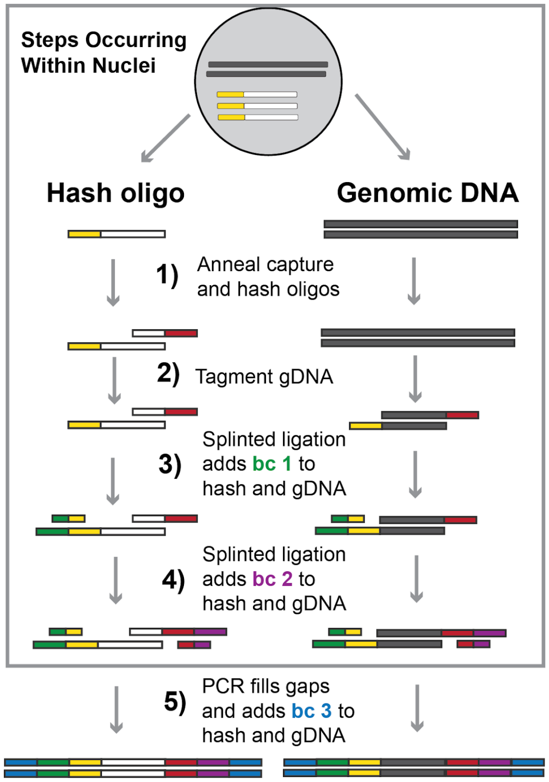


**Supplementary Figure Fig. 12:** Schematic of sciPlex-ATAC3 combinatorial indexing strategy.

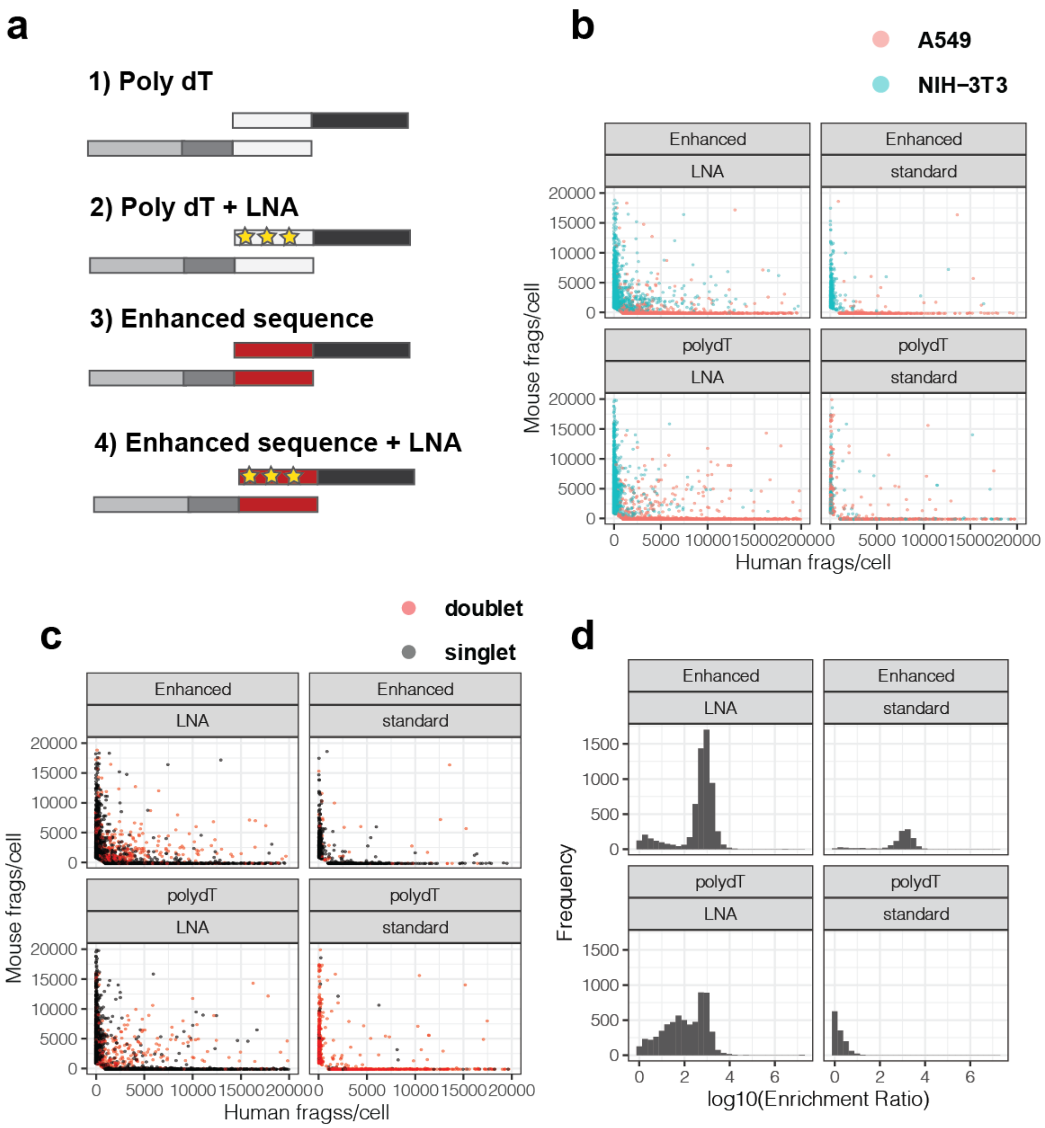


**Supplementary Figure Fig. 13:** Capture oligos with increased affinity enable hashing with ligation based combinatorial indexing. **a)** Cartoon representations of varied hash and capture oligos tested. The hash and capture oligos are represented by the bottom and top molecules, respectively. **b)** Scatter plots of unique human and mouse fragments per cell, faceted by hash and capture oligos used during library preparation. Colors show species assignments based on the dominating associated hash label for each cell. **c)** Scatter plots of human and mouse fragments per cell, faceted by library preparation conditions. Colors identify doublets assigned based on the inability to confidently assign a dominant associated hash label within a cell. **d)** Histogram of hash label enrichment scores for each nucleus, where *Enrichment Score* = *x/y*, where *x* = counts for the most common hash label within a cell and *y* = the second most common hash label within a cell. For cells with only one hash ID, the enrichment score was set as the number of hash reads (to avoid infinite values).

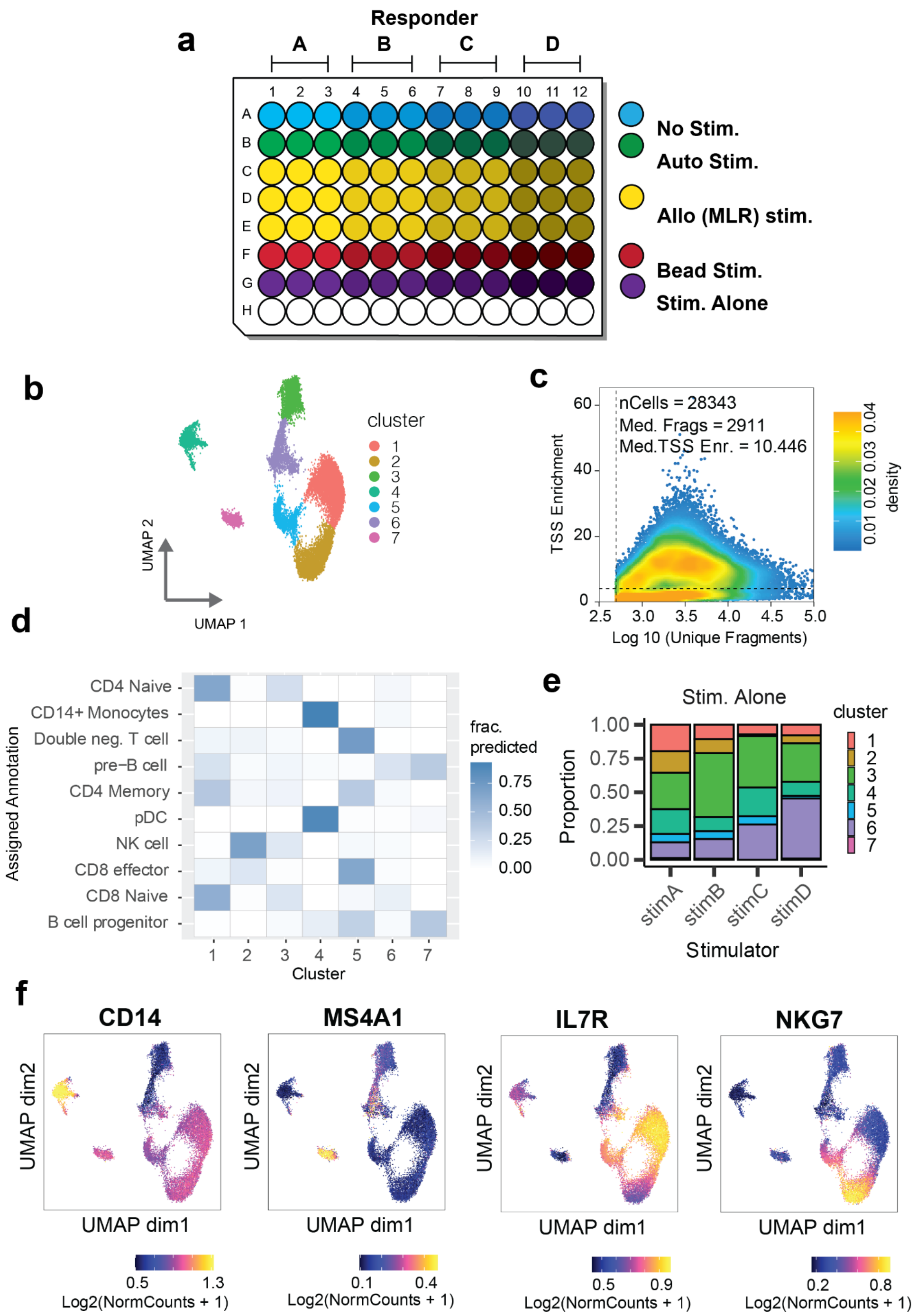


**Supplementary Figure Fig. 14:** Annotating immune cell types recovered from mixed lymphocyte reactions. **a)** Scatter plot showing the relationship between recovered fragments per cell and TSS enrichment. Dotted lines represent baseline per-cell cutoffs for each value. Cells passing these cutoffs were further filtered based on hashing (see methods). **b)** Diagram of experimental culture well layout for all conditions. **c)** UMAP representation of chromatin profiles from recovered cells colored by clusters identified with Monocle3(Cao *et al*, 2019). **d)** Heatmap depicting the fraction of cells from each cluster (shown in B) with transferred cell type assignment labels from published scRNA-seq on human PBMCs (10X genomics, pbmc_10k_v3). **e)** Proportion of cells from each stimulation-alone condition within each cluster. **f)** UMAPs colored by smoothed gene-marker accessibility scores. Gene accessibility scores were determined using ArchR and smoothing was performed with MAGIC(van Dijk *et al*, 2018), as implemented in the ArchR package(Granja *et al*, 2021).

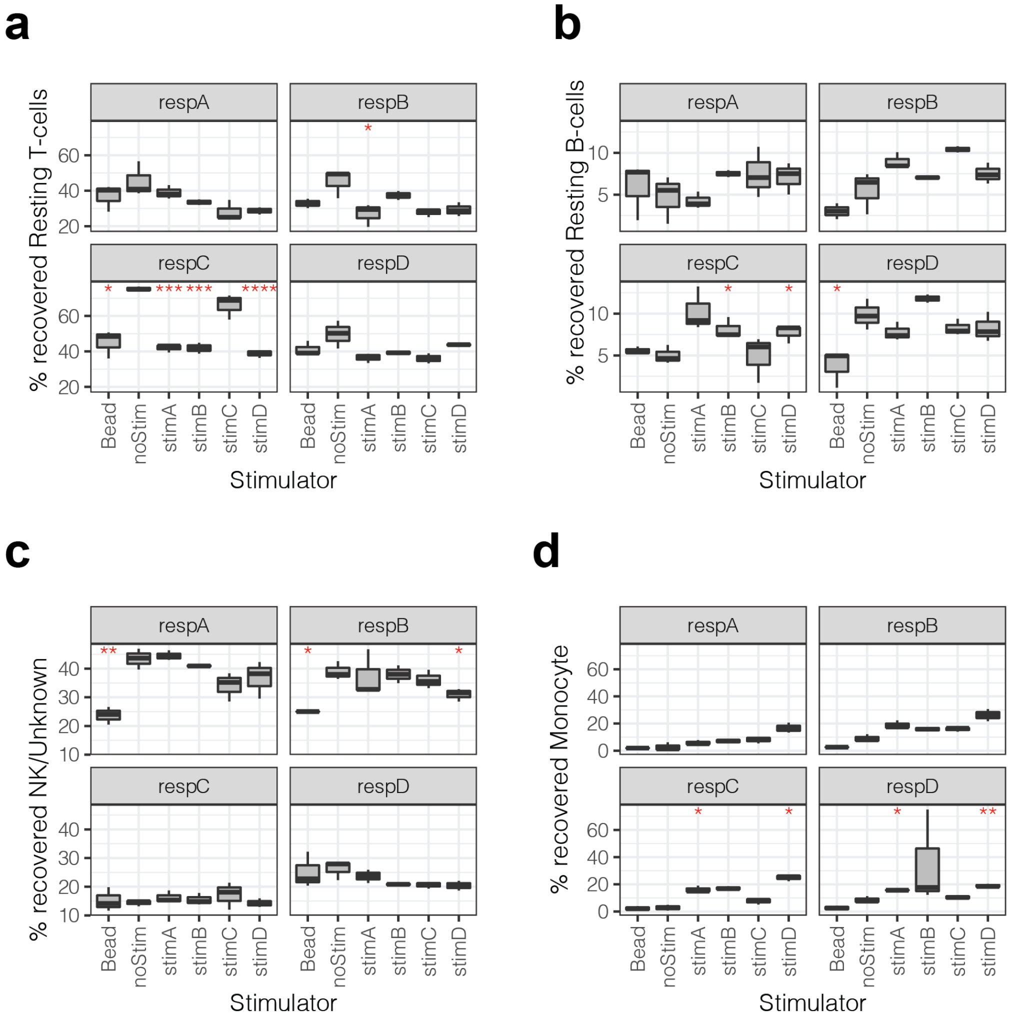


**Supplementary Figure Fig. 15:** Percentage of resting T-cells **(a),** B-cells **(b),** NK/unknown cells **(c)** Monocytes **(d)** recovered from each condition. Variation in cell type recovery was determined by separately quantifying each of three biological replicates for all conditions. Red asterisks indicate significant difference in the mean relative to no-stim for each responder (Student's t-test; *: p <0.05; **: p <0.01, ***: p < 0.001; ****: p < 0.0001).

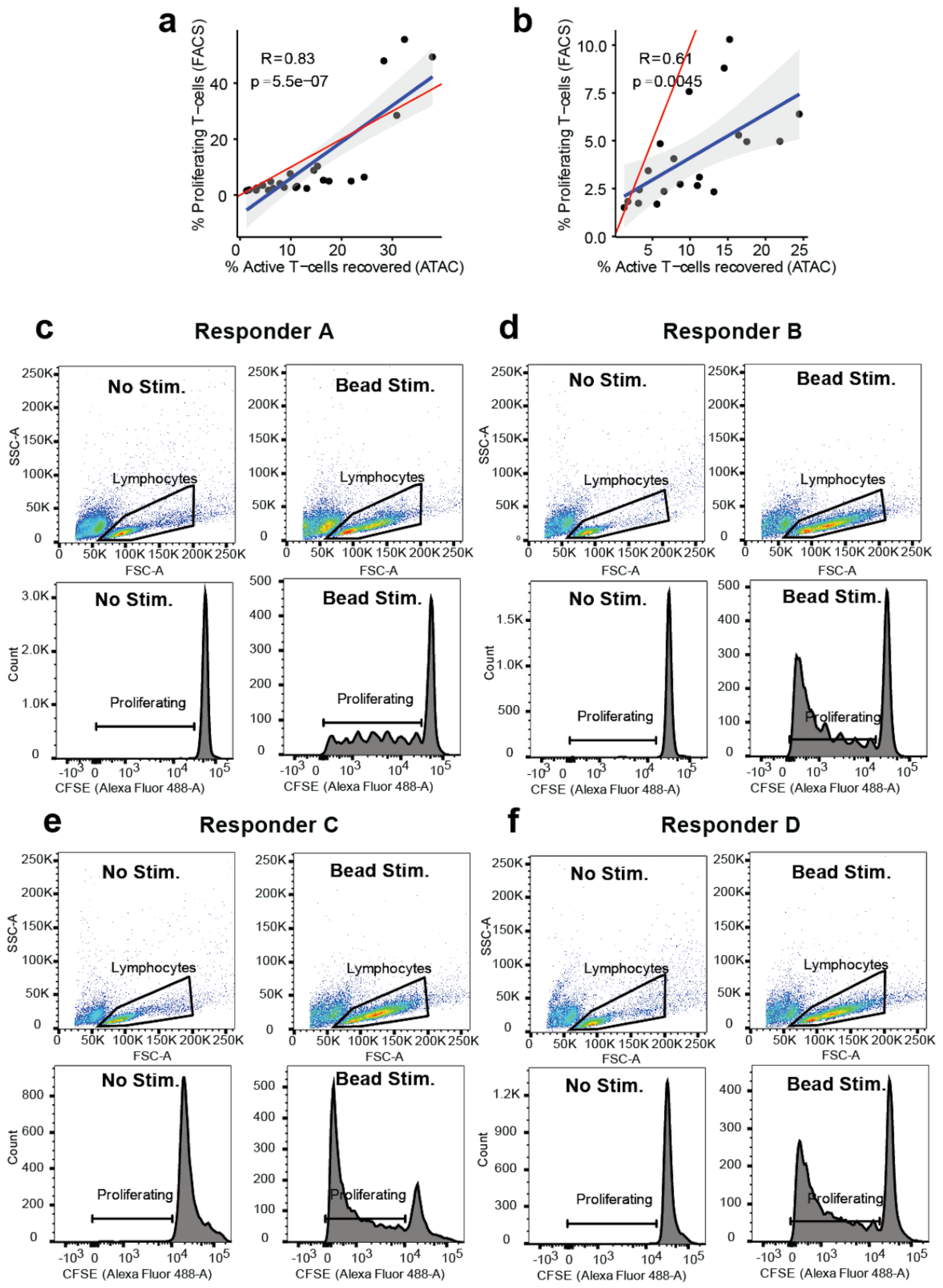


**Supplementary Figure Fig. 16:** Activated T-cell recovery with sciPlex-ATAC3 correlates well with CFSE staining. **a)** Scatter plot showing the relationship between sciPlex-ATAC3 based proportions of activated T-cells and CFSE staining-based measurements of lymphocyte proliferation across all conditions (excluding stim. alone). **b)** The same scatterplot as in a., but excluding bead-stimulated samples. **c-f)** examples of FACS populations (top) and CFSE staining (bottom) in unstimulated (left) and bead stimulated (right) conditions for each responder.

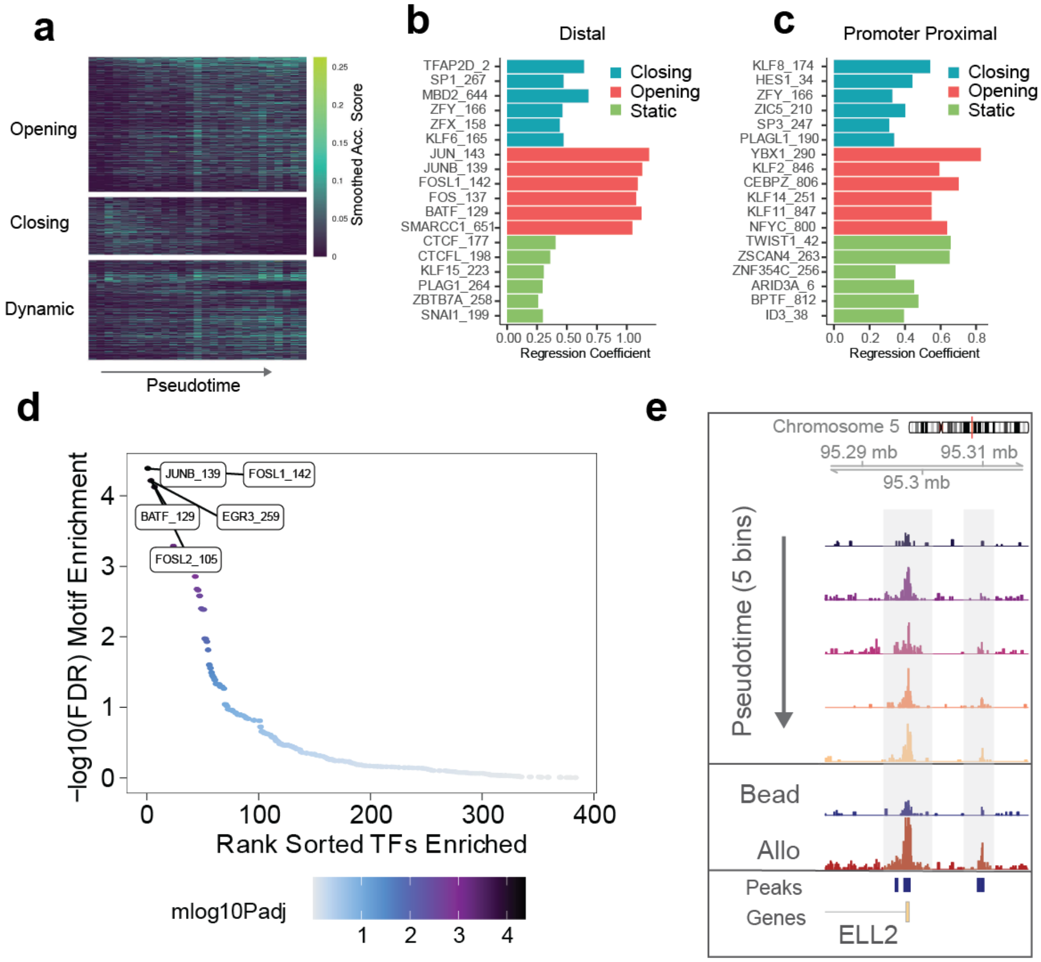


**Supplementary Figure Fig. 17:** Allogeneic stimulations elicit responses from distinct regulatory sites associated with T-cell activation and differentiation. **a)** Heatmap of raw accessibility scores (fraction cells accessible in each bin) across the pseudotime trajectory within activated T-cells only. **b&c)** Top motifs explaining whether a distal (B) or promoter-proximal (C) DA site is classified as closing (blue), opening (red) or static (non-DA, green). **d)** Ranked adjusted p-values for motif enrichment within DA sites which are open in MLR (allo)-stimulated cells compared with bead-stimulated cells. **e)** Browser tracks of pseudobulk accessibility read coverage for cells grouped into five trajectory bins (top) or by stimulation type (bottom two tracks). Y-axes are the same for all tracks (range 0-30) and represent pseudo bulk read coverage, normalized by reads within promoters for each group.

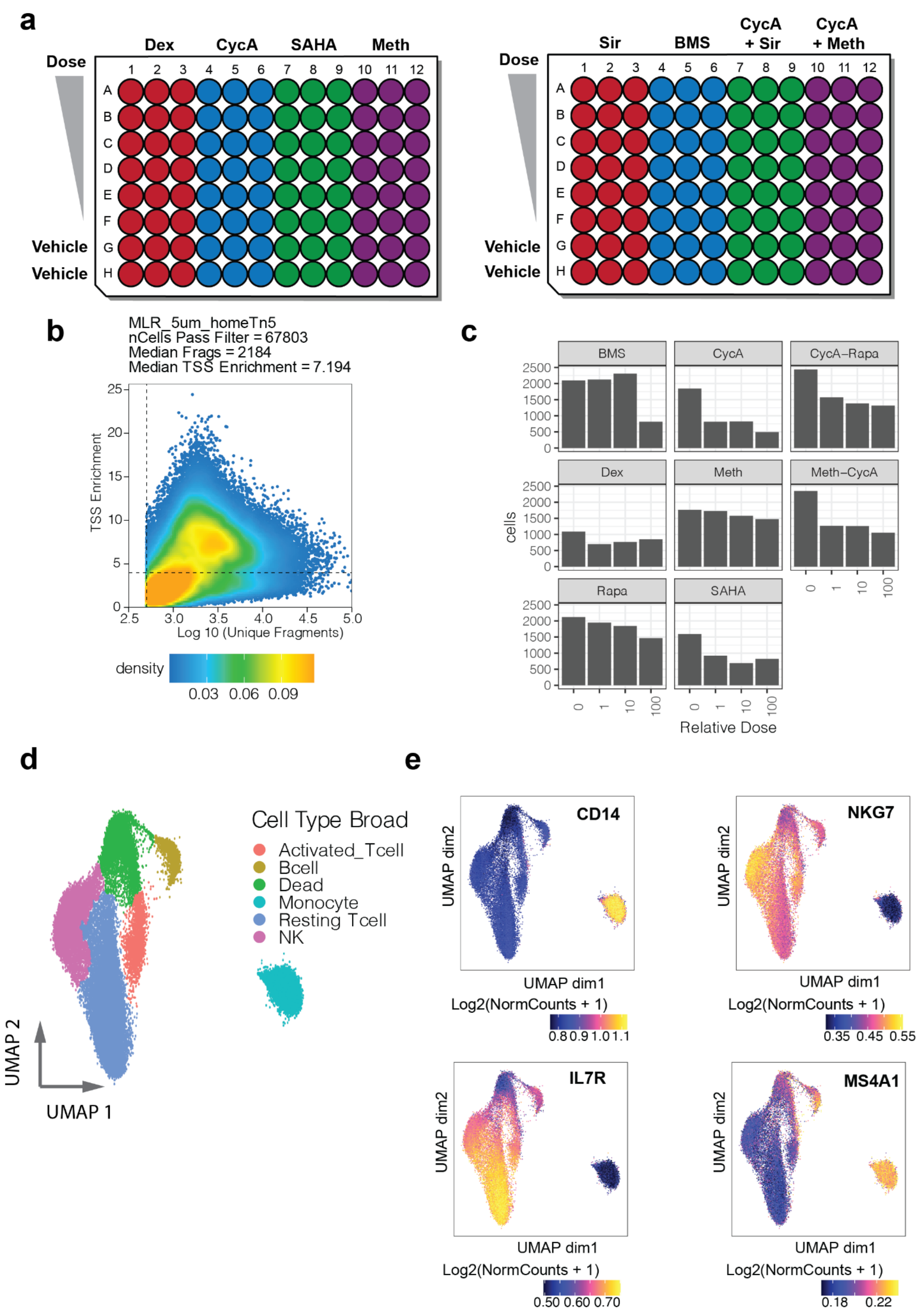


**Supplementary Figure Fig. 18:** SciPlex-ATAC3 supports combinatorial chemical perturbations within mixed lymphocyte reactions. **a)** Illustration of multi-plate experimental culture and treatment scheme. **b)** Scatter plot showing the relationship between recovered fragments per cell and TSS enrichment. Dotted lines represent baseline per-cell cutoffs for each value. Cells passing these cutoffs were further filtered based on hashing (see methods) **c)** Barplots show the number of filtered cells recovered from each treatment group. **d)** UMAP representation of chromatin profiles from recovered cells colored broad cell-type annotations (n =36,511 ). **e)** UMAPs colored by smoothed gene-marker accessibility scores. Gene accessibility scores were determined using ArchR and smoothing was performed with MAGIC(van Dijk *et al*, 2018), as implemented in the ArchR package.

| Drug | vehicle | dose_1 | dose_10 | dose_100 |
|---|---|---|---|---|
| BMS | DMSO | 0.5uM | 5uM | 50uM |
| CycA | DMSO | 0.25uM | 2.5uM | 25uM |
| CycA_Rapa | DMSO | 0.125uM, 0.25nM | 1.25uM, 2.5nM | 12.5uM, 25nM |
| Dex | Ethanol | 0.25uM | 2.5uM | 25uM |
| Meth | DMSO | 0.1uM | 1uM | 10uM |
| Meth_CycA | DMSO | 0.05uM, 0.125uM | 0.5uM, 1.25uM | 5uM, 12.5uM |
| Rapa | DMSO | 0.5uM | 5uM | 50uM |
| SAHA | DMSO | 5uM | 50uM | 500uM |

**Supplementary Table 5:** Mixed lymphocyte reaction treatment doses. For drug combinations, doses are listed in respective order.

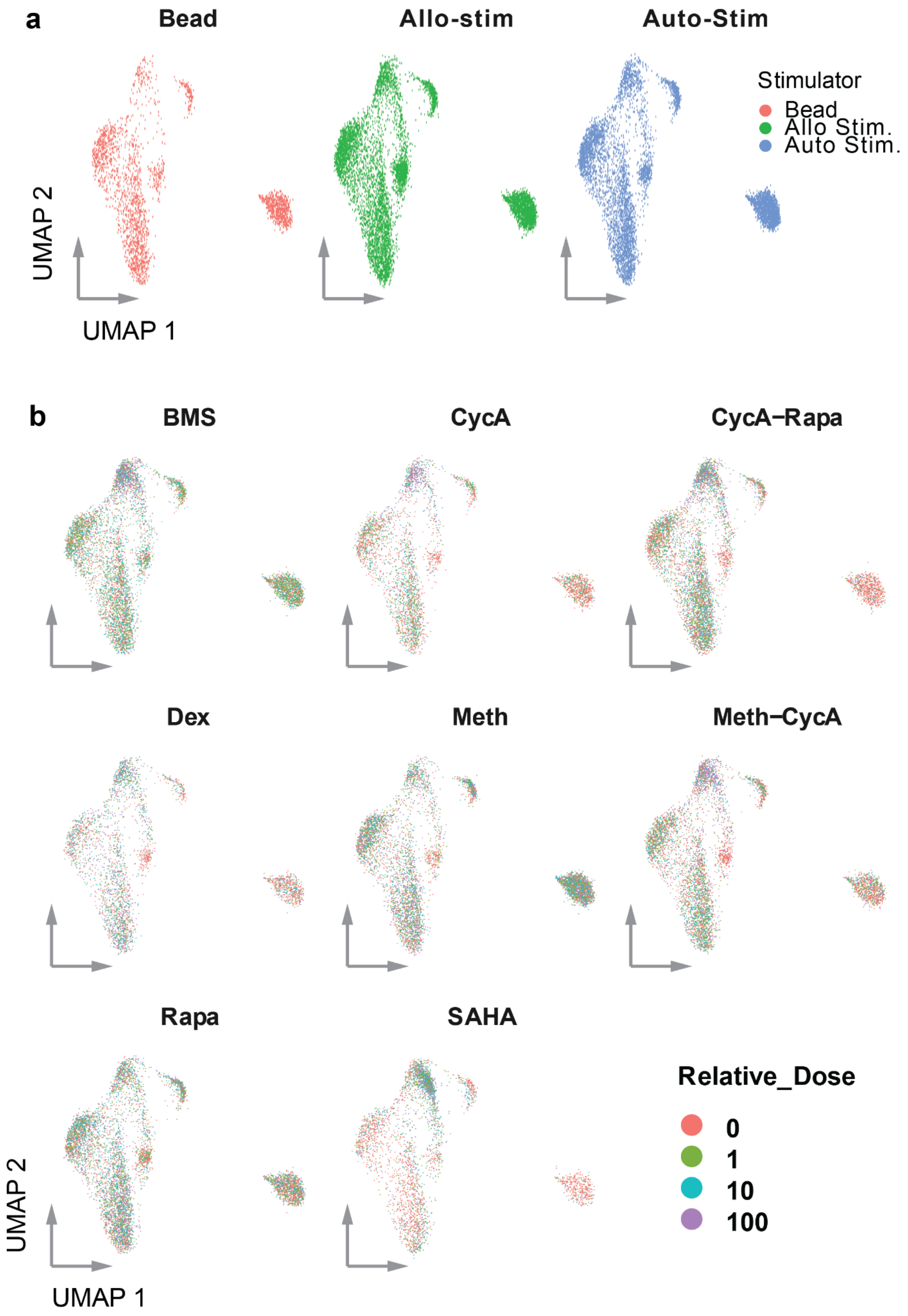


**Supplementary Figure Fig. 19:** Chemical perturbations alter the distribution of cell types and phenotypes recovered from mixed lymphocyte experiments. **a)** UMAPs of nuclei faceted by stimulation groups. **b)** UMAPs of nuclei faceted by compound/combination, with individual nuclei colored by the relative dose treatments.

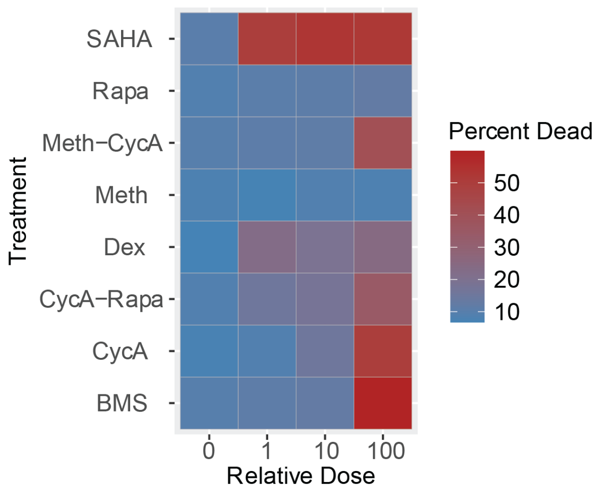


**Supplementary Figure Fig. 20:** Percent recovery of dead/irradiated cells in relation to treatment group and dose.

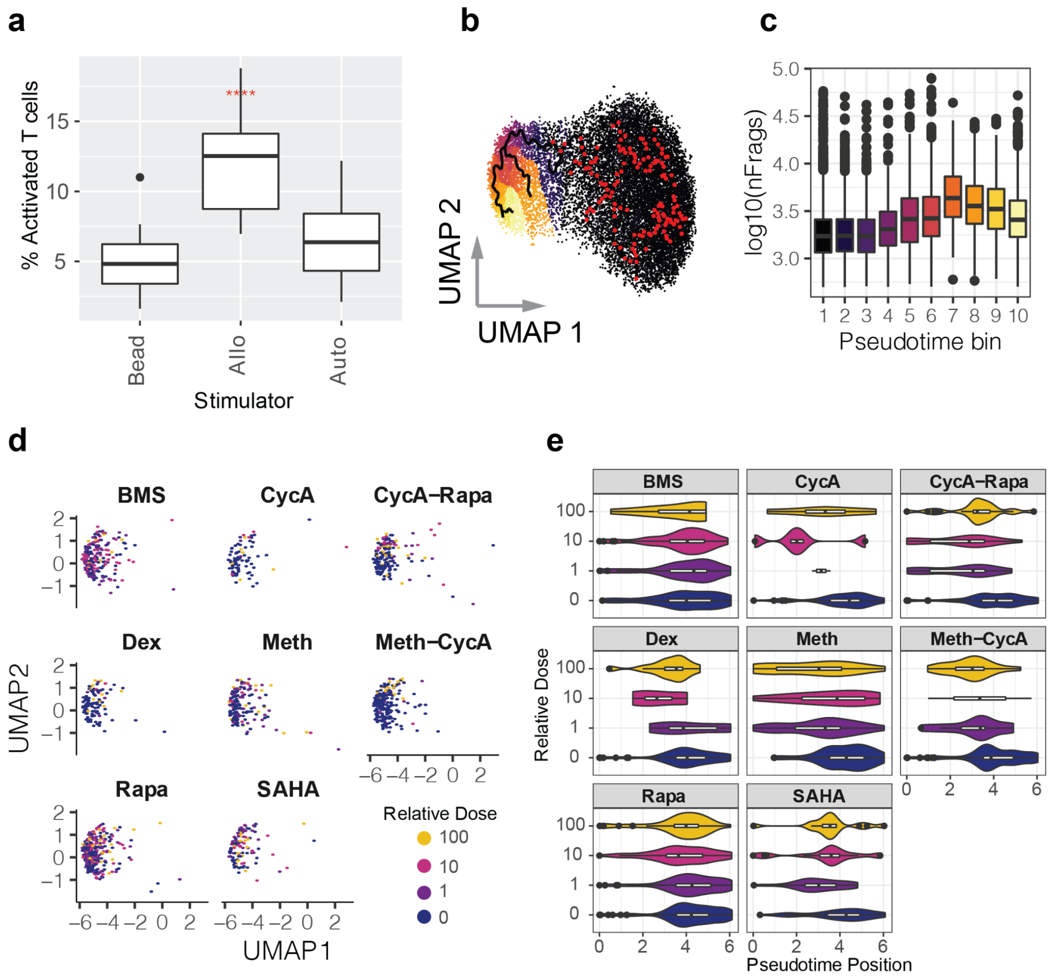


**Supplementary Figure Fig. 21:** Allo-activated T-cells are phenotypically perturbed by immunosuppressants compounds. **a)** Percentage of activated T-cells recovered from vehicle treated conditions for each stimulation condition. Variation in cell type recovery was determined by separately quantifying two vehicle treated biological replicates from each chemical treatment for the three stimulation conditions (n = 16). Red asterisks indicate significant difference in the mean relative to no-stim for each responder (Student's t-test; ****: $p < 0.0001$). See methods regarding bead stimulation conditions. **b)** UMAP representation of chromatin profiles of T-cells from the allo-stimulated wells of the MLR-drug experiment, colored by pseudotime bins as determined using the *learn_graph* function from Monocle3 and using resting T-cells as the trajectory roots. **c)** Distribution of frags per cell recovered within bins across the T-cell activation trajectory. **d)** UMAP representation of chromatin profiles from only the allo-activated T-cells faceted by compound/combination treatment and colored by relative dose. **e)** Distribution of pseudotime positions of activated T-cells from allogeneically (allo) stimulated conditions faceted by compound/combination treatment and grouped by relative dose. Figures b-e only reflect data from nuclei recovered from allo-stimulated conditions.

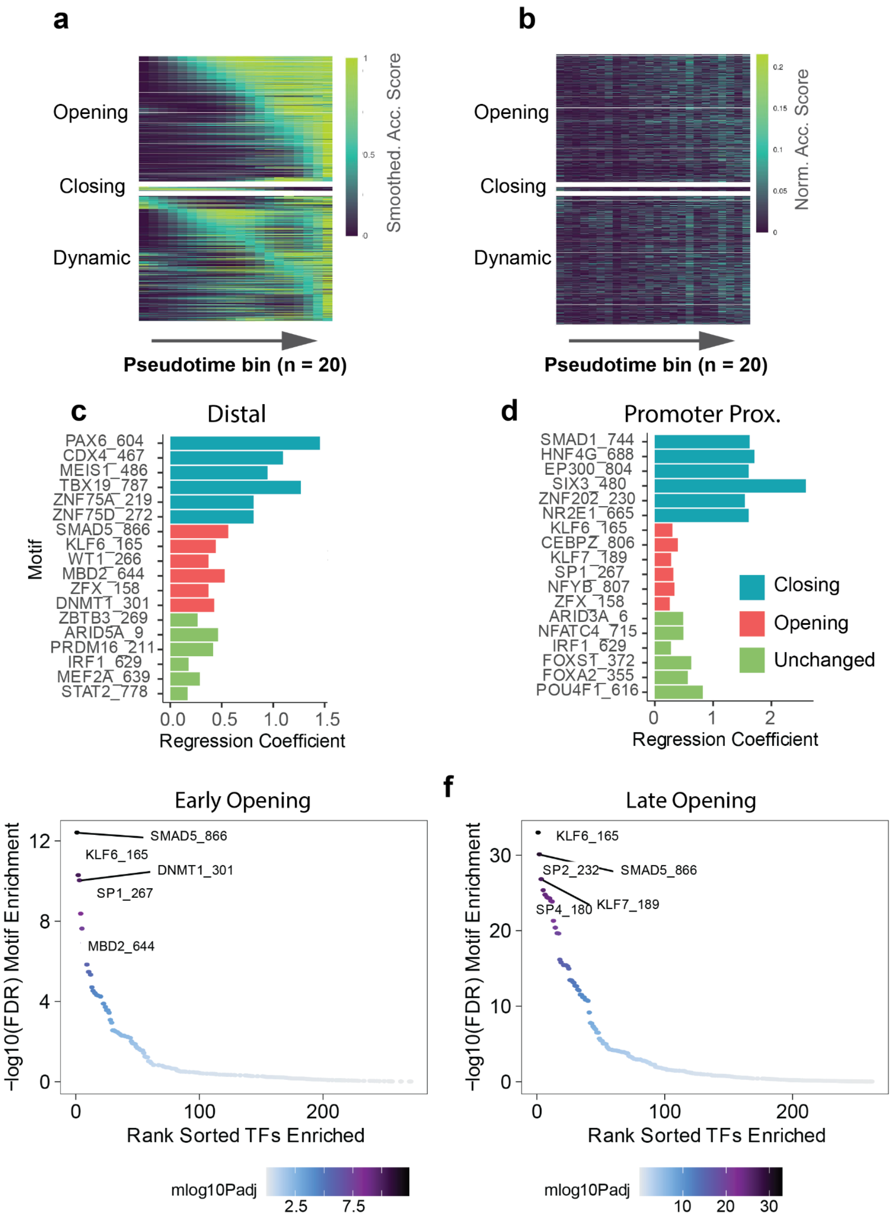


**Supplementary Figure Fig. 22:** Within allo-activated T-cells, immunosuppressant treatment primarily impedes increased accessibility. **a)** Heatmap showing smoothed accessibility scores for peaks found to change ($p < 0.05$) across the pseudotime trajectory, restricted to only allo-activated T-cells. Pseudotime bins reflect the same 20 value ranges used in figure 4c. **b)**Heatmap of raw accessibility scores (fraction cells accessible in each bin) across the pseudotime trajectory within allo-activated T-cells only. **c&d)** Top motifs explaining whether a distal (n = 592; B) or promoter-proximal (n =515; C) DA site is classified as closing (blue), opening (red) or unchanging (non-DA, green). **e&f)** Ranked adjusted p-values for motif enrichment within DA sites which open early (n = 146) (D) or late (n = 961) (E) across the trajectory within allo-activated T-cells. Early and late opening sites were defined as those in which the half maximum smoothed accessibility was found within the first 10 (of 20) pseudotime bins (early), or latter 10 bins (late).
